# Affinity purification mass spectrometry characterization of the interactome of receptor tyrosine kinase proline rich motifs in cancer

**DOI:** 10.1101/2024.05.23.595484

**Authors:** Christopher M. Jones, Arndt Rohwedder, Kin Man Suen, Safoura Zahed Mohajerani, Antonio N. Calabrese, Sabine Knipp, Mark T. Bedford, John E. Ladbury

## Abstract

Receptor tyrosine kinase (RTK) overexpression is linked to the development and progression of multiple cancers. RTKs are classically considered to initiate cytoplasmic signalling pathways via ligand-induced tyrosine phosphorylation, however recent evidence points to a second tier of signalling contingent on interactions mediated by the proline-rich motif (PRM) regions of non-activated RTKs. The presence of PRMs on the C-termini of >40% of all RTKs and the abundance of PRM-binding proteins encoded by the human genome suggests that there is likely to be a large number of previously unexplored interactions which add to the RTK intracellular interactome. Here, we explore the RTK PRM interactome and its potential significance using affinity purification mass spectrometry and in silico enrichment analyses. Peptides comprising PRM-containing C-terminal tail regions of EGFR, FGFR2 and HER2 were used as bait to affinity purify bound proteins from different cancer cell line lysates. 490 unique interactors were identified, amongst which proteins with metabolic, homeostatic and migratory functions were overrepresented. This suggests that PRMs from RTKs may sustain a diverse interactome in cancer cells. Since RTK overexpression is common in cancer RTK PRM-derived signalling may be an important, but as yet underexplored, contributor to negative cancer outcomes including resistance to kinase inhibitors.

## INTRODUCTION

Receptor tyrosine kinases (RTKs) are key mediators of intracellular signals controlling cellular growth, proliferation and motility^1^. Stimulation of transmembrane RTKs by a cognate extracellular ligand generally results in homo- or hetero-dimerization and subsequent autophosphorylation of tyrosine (pTyr) residues within cytoplasmic C-terminal RTK tails. These form docking sites for the binding of Src homology 2 (SH2), phosphotyrosine binding (PTB) and other-related domains. The development and progression of a wide range of malignancies are linked to signalling derived from RTKs^2^. To date, autophosphorylation of RTKs has been considered the predominant mediator of their oncogenic potential^2^. Despite this, therapeutic strategies to prevent RTK autophosphorylation, such as through the use of directed small molecule tyrosine kinase inhibitors, are not universally successful^3^. Further, in many cases, oncogenesis is linked to RTK protein overexpression rather than directly to increased RTK activation^2^.

Binding of proline-rich sequences by Src homology 3 (SH3) domains is critical to the assembly of a number of signalling complexes^4–13^. A comprehensive study mapping the physical and functional interactome of human RTKs identified SH3 domain-containing proteins as those most commonly bound; albeit without providing evidence for their binding sites^14^. Of the 58 RTKs encoded by the human genome, 24 feature a canonical proline-rich motif (PRM) capable of recognising SH3 domains within their cytoplasmic C-terminal tail sequence (**Fig. 1a, Supp. Table 1**)^15^. Combined with the identification of in excess of 300 sequences for SH3 domains across over 200 different proteins expressed in humans^16^ there exists to potential for a multitude of previously unstudied interactions.

**Figure 1:**
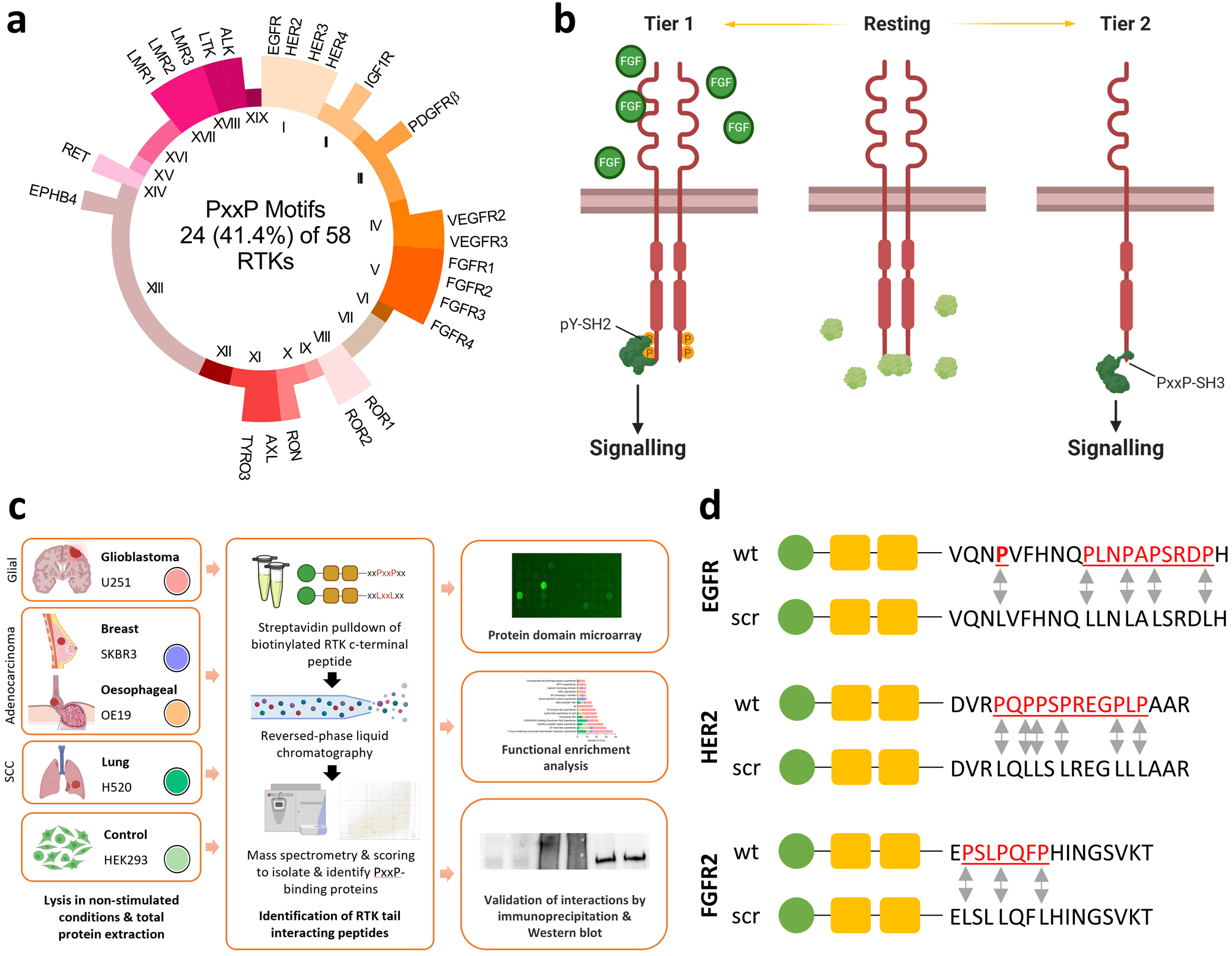
Streptavidin affinity pulldown identifies binding partners for the proline-rich C-terminal tail sequences of the receptor tyrosine kinases (RTKs) epidermal growth factor receptor (EGFR), fibroblast growth factor receptor 2 (FGFR2) and erb-B2 (ERBB2/HER2). (a) A schematic overview of the prevalence of proline rich motifs (PRMs) within each of the nineteen RTK groups encoded by the human genome. All 58 RTKs contribute to the circle equally, with only the 24 that incorporated a PRM labelled. Tail sequences are summarised in Supplementary **Table 1**. (b) A schematic of the proposed tiers of RTK-derived signalling is shown using FGFR2. The *resting* state represents conditions of low ligand availability but a relative excess of the adaptor protein growth factor receptor bound protein 2 (GRB2), which binds via SH3 domains to proline-rich sequences on the cytoplasmic tail of the RTK to form a stable heterotetramer. *Tier 1* refers to a canonical signalling mechanism through which ligand stimulation (or an activating mutation (not shown)) results in autophosphorylation of cytoplasmic C-terminal tail tyrosine residues, to which effector proteins bind via Src Homology 2 (SH2) or protein tyrosine binding (PTB) domains. *Tier 2* represents conditions of low ligand availability but a relative excess of RTK compared with GRB2, such as may occur following RTK protein overexpression in instances of gene amplification. In these conditions, binding of effector proteins to RTK PRMs via SH3 and related domains results in downstream signalling. (c) A summary of the experimental approach used to identify binding partners of PRM-containing tail regions of EGFR, FGFR2 and HER2 in cells representing glioblastoma (U251), lung squamous cell carcinoma (H520), oesophageal adenocarcinoma (OE19), breast adenocarcinoma (SK-BR-3) or a control cell line (HEK293T) that stably overexpresses FGFR2. Lysates from each cell line were individually incubated with bait peptides representing PRM-containing wild-type RTK c-terminal tail sequences or scrambled control sequences in which proline residues were replaced with leucine and which were devoid of tyrosine residues. Mass-spectrometry was used to identify bound peptides, which were identified using MaxQuant and compared across bait and control tail sequences using SAINTexpress and Perseus. Functional enrichment of identified interactors was used to characterise their structure and function. A subset of interactors were evaluated through live cell imaging to confirm their proximity and by gene knockdown analysis in order to assay their function in conditions in conditions of low RTK phosphorylation. (d) Bait peptide sequences used for the streptavidin pulldown. Tail sequences of EGFR (residues 1105-1124), HER2 (residues 1141-1161) and FGFR2 (residues 806-821) were covalently bound to biotin via two polyethylene glycoPEG) spacers. Proline residues and PRMs are underlined and highlighted in red. Arrows are used to signify points at which proline residues in the wild-type (wt) bait polypeptide were replaced by leucine residues within scrambled (scr) control bait polypeptide for each of the three RTK sequences.

Recent evidence to indicate that the recruitment of signalling proteins to these sites, in the absence of RTK upregulation, associates with signalling activity and pathological outcomes, but that the extent to which this defines cancer outcomes is unclear. A notable exception is the known interaction of the C-terminal SH3 domains of the adaptor protein growth factor receptor bound protein 2 (GRB2) and the phospholipase Cγ1 (PLCγ1) with a PRM within the C-terminal tail of fibroblast growth factor receptor 2 (FGFR2)^17,18^. In the absence of RTK activity, intracellular, concentration-dependent competition between the SH3 domains of GRB2 and PLCγ1 for the receptor PRM regulates activity of the phospholipase and AKT-mediated cell proliferation and motility^19,20^. There is, in addition, evidence for SH3 domain-mediated binding of the proto-oncogene protein tyrosine kinase, FYN, to a cytoplasmic C-terminal tail PRM within the ERBB2 (HER2) receptor tyrosine kinase^21^. Other domains are also recognised to bind PRMs but have not yet been explored in the context of RTK PRM sequences. These include WW domains, Ena/Vasp homology domain 1 (EVH1) domains, glycine-tyrosine-phenylalanine (GYF) domains, ubiquitin E2 variant (UEV) domains and single-domain profilin proteins^22^.

Together, these findings suggest the presence of a second tier of RTK-derived signalling that is not contingent on ‘on/off’ ligand-induced pTyr-mediated signalling, but on interactions that occur in the absence of ligand stimulation by C-terminal PRMs (**Fig. 1b**). Crucially, this signalling (henceforth termed Tier 2) is dependent on the relative intracellular concentration of cognate effector proteins. This means that conditions that drive fluctuations in concentration of these proteins (e.g., environmental stress) will permit proteins to prevail in interactions with specific RTKs and initiate different patterns of signalling. Within the context of cancer, most RTK-related research has been focused on activating mutations that hyperstimulate pTyr-mediated Tier 1 signalling^2^. There is, however, accumulating evidence that a number of cancers are characterised, and their outcomes, at least in part, dictated by frequent protein overexpression of RTKs^2^. This occurs through diverse processes including genomic amplification, loss of negative regulation and the increased transcription and translation of RTK-encoding genes. The result is a significant increase within a cell in the local concentration of overexpressed RTKs, many of which harbour PRMs capable of mediating non-canonical Tier 2 signalling.

Despite the potential importance of this to cancer outcomes, the pathways through which RTK PRMs mediate signalling are not known. Given this, we sought to uncover and characterise the PRM interactome of the RTKs epidermal growth factor receptor (EGFR), FGFR2 and ERBB2/HER2 in cell lines resembling oesophageal adenocarcinoma (OAC), breast adenocarcinoma (BrAC), glioblastoma (GBM) and lung squamous cell carcinoma (LSCC). These are malignancies in which amplification or overexpression of the three chosen RTKs is frequently identified and has been shown to be associated with survival outcomes^2,23–32^. In studying these, we provide evidence for a diverse RTK PRM interactome that is enriched for metabolic, homeostatic and pro-migratory signalling pathways. We also provide further evidence for the importance to cancer outcomes of SH3-mediated interactions with the PRM of RTKs in the absence of pTyr upregulation.

## RESULTS

### The RTK PRM-containing tail region interactome across RTKs and cancer cell lines

In order to uncover the interactomes for specific RTK PRMs, a PRM-incorporating, tyrosine depleted, C-terminal tail regions from each of the RTKs; EGFR, FGFR2 and ERBB2/HER2 was used as bait to affinity purify bound proteins in cell lines resembling OAC, BrAC, GBM and LSCC as well as a non-cancerous cell line (HEK293T); as summarised in **Fig. 1c**. Captured proteins were compared to those bound to similar bait peptides in which PRMs were replaced by leucine residues (**Fig. 1d**). The inclusion of leucine residues precludes the PRM from adopting the canonical PPII helical structure required for ligand recognition. PRM-interacting proteins were subsequently identified by high-resolution mass spectrometry.

Using the probabilistic SAINTexpress scoring algorithm, a total of 490 unique proteins were identified as interactors for at least one RTK PRM-containing tail region in at least one of the studied cell lines (**Fig. 2a, Supp. Tables 2-17**) (see Methods). Across all five studied cell lines, the largest number of interactors was seen with the EGFR PRM-tail region (n=454). In contrast, 155 interactors were identified for the FGFR2 PRM tail region and only two for the HER2 PRM tail region. For the EGFR PRM-tail region, the greatest number of interactors were seen for U251 cells (n=294), with 67 identified for H520 cells, 48 for OE19 cells and 29 for SKBR3 cells. For the FGFR2 PRM-tail region, 92 interactors were identified for SKBR3 cells, 28 for H520 cells, 24 for U251 cells and ten for OE19 cells. The identified number of interactors was lowest in HEK293T cells for both the EGFR (n=16) and the FGFR2 (n=1) PRM-tail regions. This discrepancy in the number of interactors identified for each cell lysate is consistent with the different expression profiles of PRM-binding proteins which presents a unique repertoire and of signalling proteins at distinct concentrations available that each PRM can accommodate.

**Figure 2:**
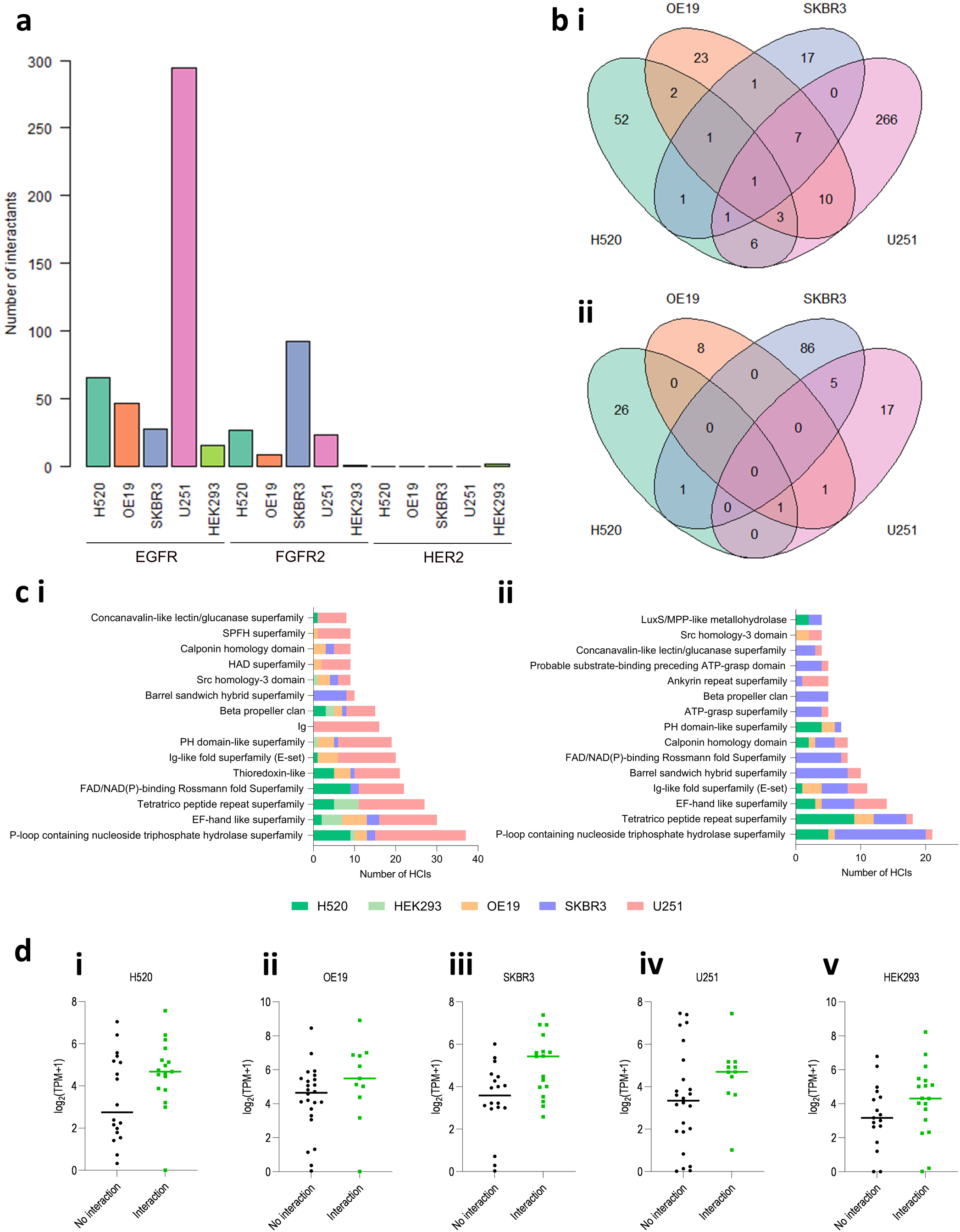
A summary of the number and structural composition of interactors for proline rich C-terminal sequences of the receptor tyrosine kinases (RTKs) epidermal growth factor receptor (EGFR), fibroblast growth factor receptor 2 (FGFR2) and erb-B2 (ERBB2/HER2). **(a)** The total number of interactors for each streptavidin pulldown experiment using a proline-rich C-terminal RTK tail sequence is shown for each of the studied cell lines: lung squamous cell carcinoma (H520), oesophageal adenocarcinoma (OE19), breast adenocarcinoma (SKBR3), glioblastoma (U251) and human embryonic kidney (HEK293T). **(b)** Venn diagrams illustrating the number of interacting proteins for the **(i)** EGFR and **(ii)** FGFR2 proline rich C-terminal tail sequences that were shared across each of the four studied cell lines. There were no shared interactors for the HER2 C-terminal tail sequence. **(c)** A summary of the most commonly identified Pfam protein clans for the **(i)** EGFR and **(ii)** FGFR2 proline rich C-terminal tail sequences and their relationship to the studied cell lines. **(d)** mRNA expression data were extracted for each studied cell line from the Cancer Cell Line Encyclopaedia for the cohort of SH3-domain containing interactors shown in **Table 2**. Expression data, shown as a pseudo-count of log2(transcripts per million (TPM)+1), are categorised by the interaction status of the protein to which they correspond. Genes corresponding to proteins not identified as an interactor for any of the studied RTKs in each specific cell line are categorised as ‘No interaction’ and genes corresponding to proteins identified as an interactor for at least one of the studied RTKs in each cell line are listed as ‘Interaction’. Data are shown for each of **(i)** H520, **(ii)** OE19, **(iii)** SKBR3, **(iv)** U251 and **(v)** HEK293T cell lines.

In total, sixty proteins bound both the EGFR and the FGFR2 proline-rich motifs: 28 (46.7%) within the same cell lines (**Supp. Table 2**). One interactor (triosephosphate isomerase, TPIS) was identified as bound to EGFR in each of the cancerous cell lines, whereas there was no consistently identified interactor for FGFR2 or HER2 (**Fig. 2b**). Thirty-three proteins were interactors for the EGFR PRM in more than one cell line, whereas three proteins were interactors for the FGFR2 and none for HER2 PRMs in more than one cell line. The two interacting proteins for HER2 (Treacle protein, TCOF; Dedicator of cytokinesis protein 7, DOCK7) - both of which were identified in HEK293T cells - were not identified as potential interactors for either EGFR or FGFR2. Across the studied RTKs and cell lines, the most frequently identified protein was the cytoskeletal protein Spectrin alpha chain, non-erythrocytic 1 (SPTAN1), which bound both the EGFR and FGFR2 PRM-tail regions in OE19 and U251 cells, in addition to the EGFR PRM-tail region in SKBR3 cells.

We characterised the protein interactors of each studied RTK PRM by classifying them by Pfam clan (**Fig. 2c, Supp. Table 18**). The P-loop-containing nucleoside triphosphate hydrolase superfamily, tetratrico peptide repeat superfamily and EF-hand like superfamily were the most frequently identified amongst the EGFR and FGFR2 Interactors. The tetratrico peptide repeat superfamily was also represented by one of the two HER2 Interactors.

We also sought to evaluate whether the expression level of identified interactors influences the probability of their identification as a cell interactor. To do so, mRNA expression of genes encoding the full list of identified RTK PRM interactors **(Supp. Table 2)** across all RTKs and cell lines was obtained for each cell line from The Cancer Cell Line Encyclopedia (CCLE)^33^. These are correlated against interaction status for each cell line in **Fig. 2d**, such that counts for genes encoding proteins not identified as an interactor for any of the studied RTKs in each cell line are grouped as ‘No interaction’ and counts for genes encoding proteins identified as an interactor for at least one of the RTKs in each cell line are grouped as ‘Interaction’. A higher median expression score was seen in each cell line for interactors, suggesting that relative protein concentration influences the probability of a PRM-SH3 interaction occurring.

These data reveal that PRMs from a subset of RTKs are able to interact with a large and diverse range of proteins from cancer cell lysates. If this PRM-mediated interactome is replicated in vivo this represents a substantial and previously over-looked signal regulating capability.

### Metabolic, homeostatic and migratory processes are overrepresented amongst interactors of RTK PRM regions

To evaluate the functional significance of Tier 2 signalling derived from RTK PRMs we identified overrepresented protein class, biological process and molecular function terms amongst interactors for the three RTK-cell line combinations demonstrating the highest number of interactors: the EGFR C-terminal tail in H520 LSCC cells and U251 GBM cells, and the FGFR2 C-terminal tail in SKBR3 BrAC cells.

The largest proportion of interactors across each of the studied RTK-cell line combinations were classed as metabolite interconversion or protein modifying enzymes (**Fig. 3a**). Other commonly identified interactors included translational and cytoskeletal proteins, scaffold/adaptor and chaperone proteins, and cytoskeletal proteins. Accordingly, amongst the interactors, the most overrepresented GO biological processes related to metabolism (‘metabolic process’), homeostasis (‘biological regulation’, ‘response to stimulus’) and cellular movement (‘localisation’, ‘locomotion’ and ‘biological adhesion’); as summarised in **Fig. 3b**. In keeping with an ability to transduce signalling, ‘catalytic activity’ was the most overrepresented GO molecular function term across each three RTK-cell line combinations (**Fig. 3c**).

**Figure 3:**
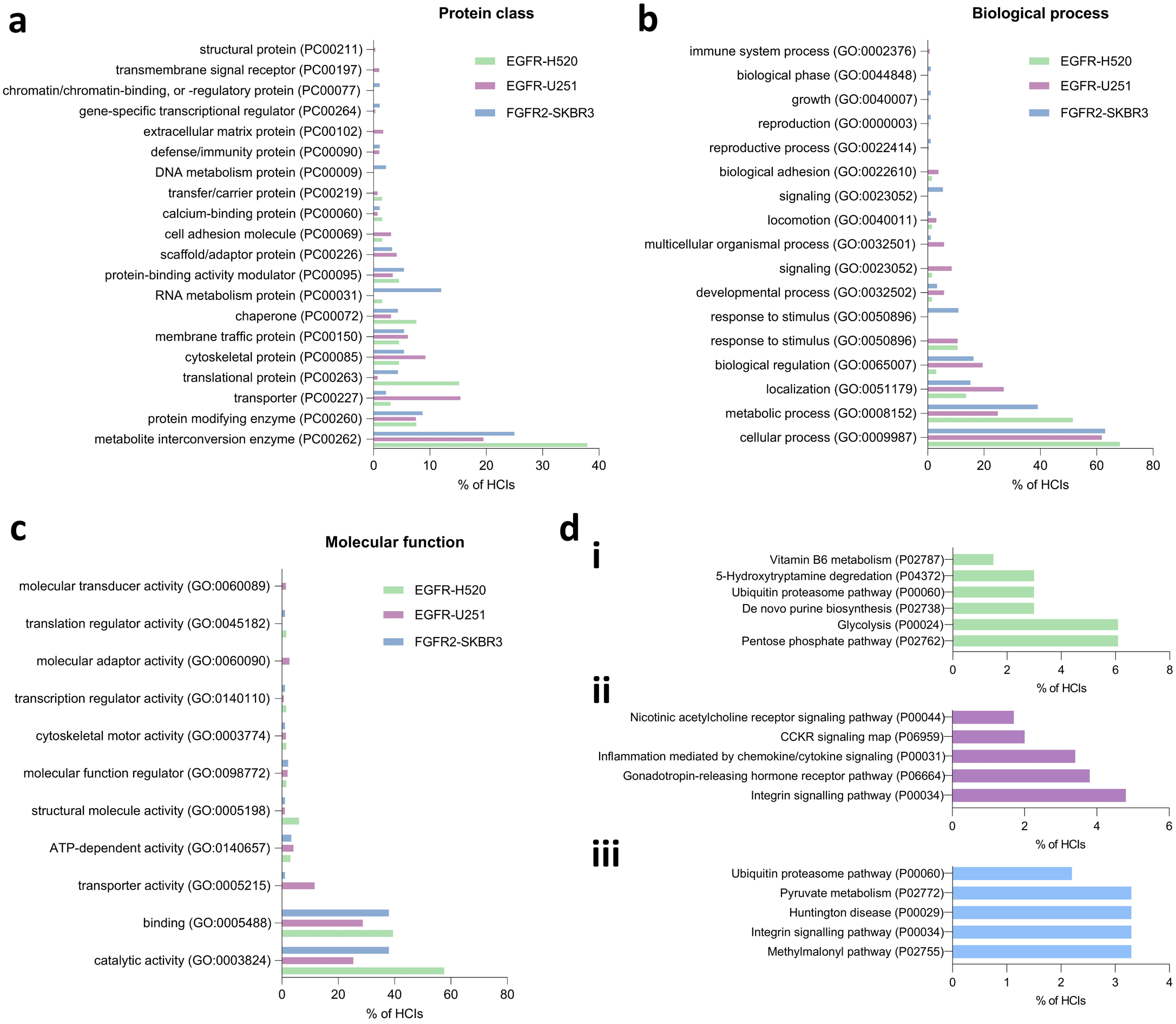
Functional characterisation of interactors of the epidermal growth factor receptor (EGFR) and fibroblast growth factor receptor 2 (FGFR2) C-terminal tail proline rich motifs (PRMs) in H520 lung squamous cell carcinoma, U251 glioblastoma and SKBR3 breast adenocarcinoma cells. **(a)** Overrepresented protein class terms amongst interactors for EGFR and FGFR2 in H520, U251 and SKBR3 cells. **(b)** Overrepresented biological process terms amongst interactors for EGFR and FGFR2 in H520, U251 and SKBR3 cells. **(c)** Overrepresented molecular function terms amongst interactors for EGFR and FGFR2 in H520, U251 and SKBR3 cells. **(d)** The top 5 overrepresented signalling pathways for interactors of **(i)** the EGFR C-terminal tail PRM in H520 cells, **(ii)** the EGFR C-terminal tail PRM in U251 cells, and **(iii)** the FGFR2 C-terminal tail PRM in SKBR3 cells. In all cases, the proportion of interactors contributing to overrepresentation of each term is shown on the x-axis.

The specific pathways overrepresented by interactors were mostly in keeping with these broad processes (**Fig. 3di-iii**). This includes metabolic pathways such as those relating to glycolysis, pyruvate metabolism and the Kreb’s cycle (methylmalonyl pathway), as well as migration-related pathways such as those relating to integrin signalling. Interestingly, in keeping with overrepresentation of immune system processes amongst interactors of the EGFR PRM in U251 GBM cells, enriched pathways amongst these interactors included ‘inflammation mediated by chemokine/cytokine signalling’ and cholecystokinin receptor ‘CCKR’ signalling (**Fig. 3dii**).

### Analysis of Tier 2 signalling mediator interactions with RTK PRM regions

Since they provide recognition sequences for a range of different protein domains, we analysed interactors for the WW, EVH1, GYF, UEV and profilin domains (**Table 1**) that are known to interact with PRMs; albeit not in the context of an RTK C-terminal tail. Nineteen EGFR interactors and seven FGFR2 interactors contained an EVH1 domain, as represented by the PH domain-like superfamily. A further four interactors of EGFR featured a profilin domain, as represented by the profilin-like superfamily. There were no recognised WW, GYF or UEV domains amongst the identified interactors.

**Table 1:**
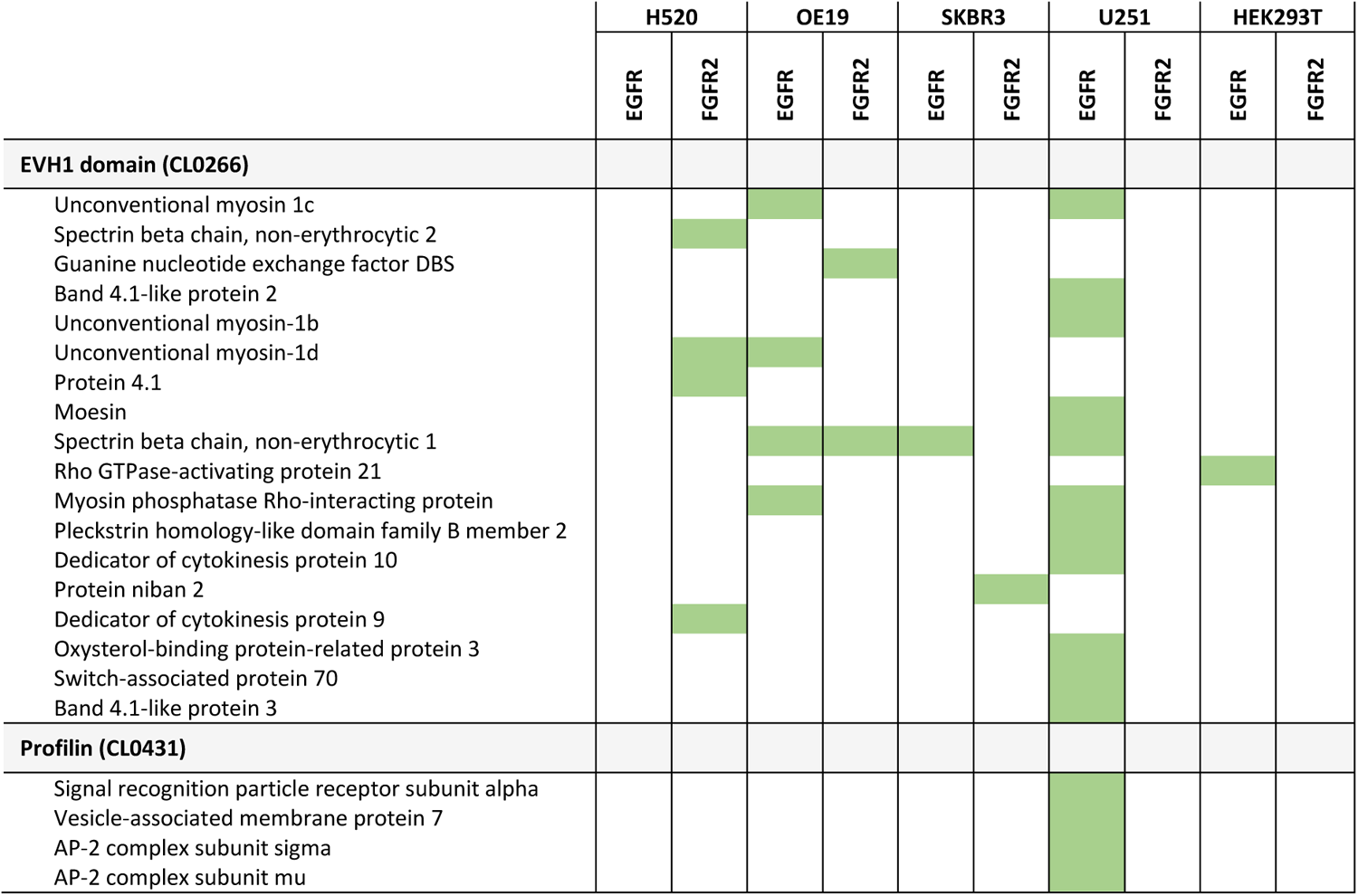
A summary of identified proline rich motif (PRM) interactors that incorporate a EVH1 or profilin domain. The EVH1 and profilin domains have previously been reported to mediate binding to PRMs. Highlighted green cells indicate the presence of a detected interaction between the named protein, listed by the relevant incorporated domain, and the C-terminal tail region of either epidermal growth factor receptor (EGFR) or fibroblast growth factor receptor 2 (FGFR2) in lysates from cells resembling glioblastoma (U251), adenocarcinoma of the lung (H520), oesophagus (OE19) or breast (SKBR3), or from the non-malignant HEK293T cell line. Interactors were identified using SAINTexpress (see Methods).

We also sought to characterise interactions between RTK PRMs and SH3 domain-containing proteins (**Table 2**). This was of particular interest given previous evidence for a role for SH3-PRM interactions in mediating deleterious cancer outcomes^19,20^. From our data, only six SH3-containing proteins were identified using SAINTexpress as interactors for at least one RTK PRM-containing tail region in at least one of the studied cell lines. This low number of reflects that interactions between PRMs and SH3 domains (K_d_ = 1-100μM) are at least ten-fold weaker than those of other RTK C-termini interactions (e.g., pTyr sites with SH2 or PTB domains)^34^. The lower affinity of the interactions does not preclude their physiological importance in signalling because, as stated above, the interactions are equilibrium-based and hence dependent on respective concentrations of binding partners.

**Table 2:**
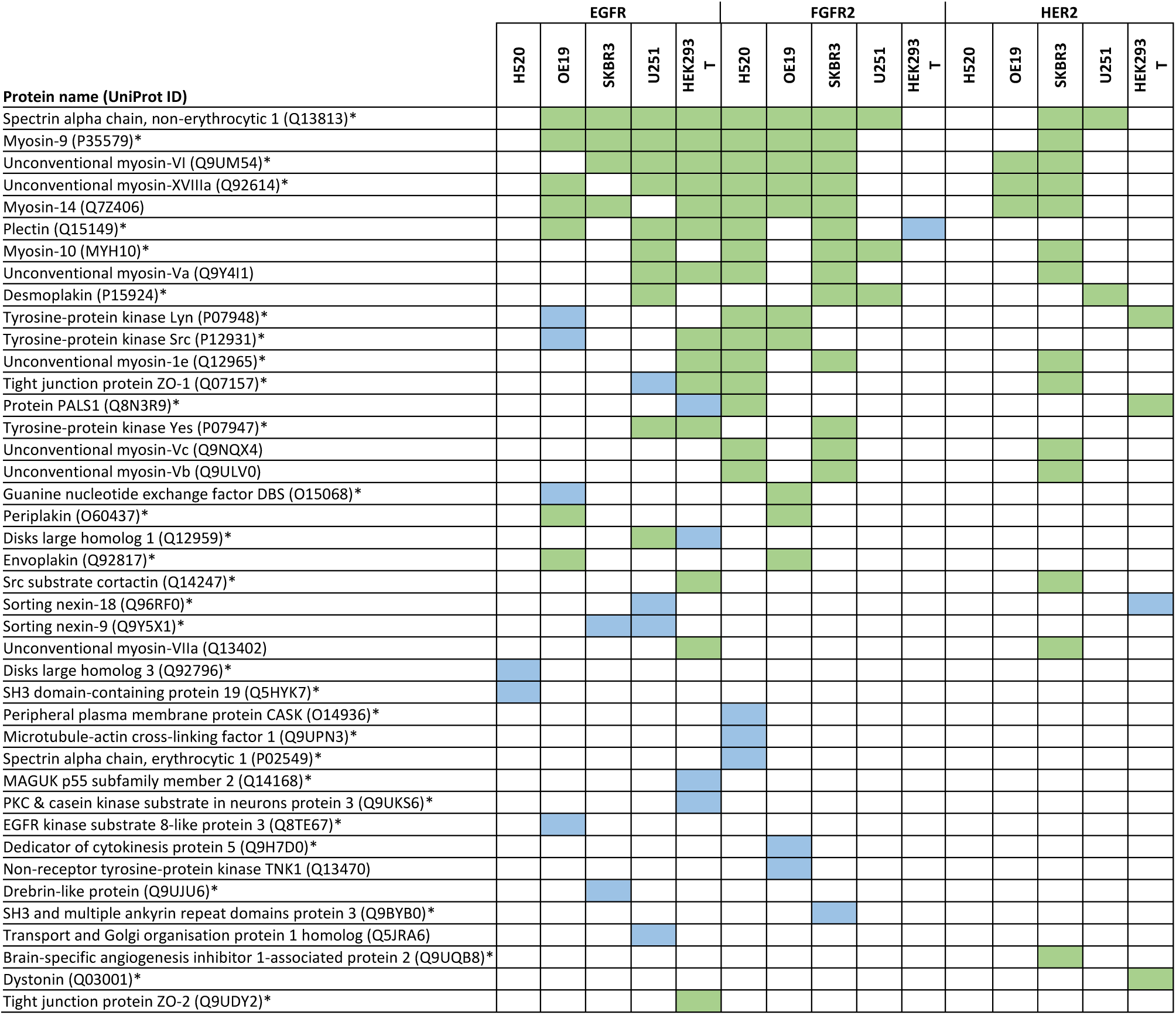
A summary of SH3 domain-containing interactors (green) and additional low confidence interactors (LCIs; blue) for each of epidermal growth factor receptor (EGFR), fibroblast growth factor receptor 2 (FGFR2) and erb-B2 (ERBB2/HER2) in cell lines representing squamous cell lung cancer (H520), oesophageal adenocarcinoma (OE19), breast adenocarcinoma (SKBR3), glioblastoma (U251) and in a control HEK293T cell line. Interactors (green) were identified using SAINTexpress and Perseus, with conventional cut off scores applied. Additional LCIs (blue) were identified by SAINTexpress using less stringent cut-off values. Proteins labelled with the GO cellular component identifier ‘plasma membrane’ are highlighted with an asterix.

Importantly, it should also be appreciated that interactions of SH3 domains are typically at least an order of magnitude weaker affinity than interactions usually identified using the AP-MS approach. Consequently, SH3 domain-containing proteins may be underrepresented amongst interactors as determined by the probabilistic SAINTexpress scoring algorithm. Since there is an extensive literature characterising individual interactions of intracellular PRMs and SH3 domains, adopting the stringency level imposed by the SAINTexpress method would deny the existence of these interactions. The data presented here support this, with the Src family tyrosine kinases SRC, LYN and YES proteins not classified as interactors by SAINTexpress despite substantial previous characterisation of their interaction with PRMs^35,36^ (**Table 2**).

Given this, we further scrutinised our data using a two-pronged approach to identify additional SH3-domain containing protein interactors: (1) using label-free quantitation of on MS-intensity based data (rather than spectral counts as implemented in SAINTexpress) and (2) using less-stringent SAINTexpress cut-off values that were tuned to allow identification of interactors that had been previously experimentally validated (see Methods). Using this approach SRC, LYN and YES were all identified as potential interactors, consistent with previous experimental data. To distinguish identification of interactors using less-stringent SAINTexpress cut-off values from the previously described data, we term these low confidence interactors (LCIs).

We explored this expanded list of interactors to identify SH3-containing proteins and were able to identify a combined total of 41 SH3-containing interactors for at least one of the RTKs in one of the studied cell lines (**Table 2**). Importantly, of the 122 observed pairwise PRM-SH3 interactions, only 26 (21.3%) were LCIs. The remaining 96 (78.7%) were identified using conventional SAINTexpress (n=5; 4.1%) or Perseus (n=81; 66.4%) scores, with five (4.1%) interactors identified by both scoring systems and a further five (4.1%) identified as interactors by Perseus but only as LCIs by SAINTexpress. This points to a potential greater sensitivity for low affinity reactions for Perseus over SAINTexpress^37^.

The median number of interactors identified across the studied cell lines was 7 (range 0-14) for EGFR and 10 (range 0-14) for FGFR2, compared with 3 (range 0-14) for HER2. Given the relatively smaller number of interactors and LCIs identified for HER2, we sought to mitigate against any impact from the screening approach by undertaking an orthogonal approach using recombinantly expressed SH3 domains immobilised on a chip and monitoring binding to a fluorescently labelled HER2 (**Fig. 4a**, control data in **Supp. Fig. 1**). This identified additional interactions with FYN (which is a Src family kinase with high sequence homology with LYN which was previously characterised^21^) and PLCƳ1 but no other assessed SH3 domain-containing proteins.

**Figure 4:**
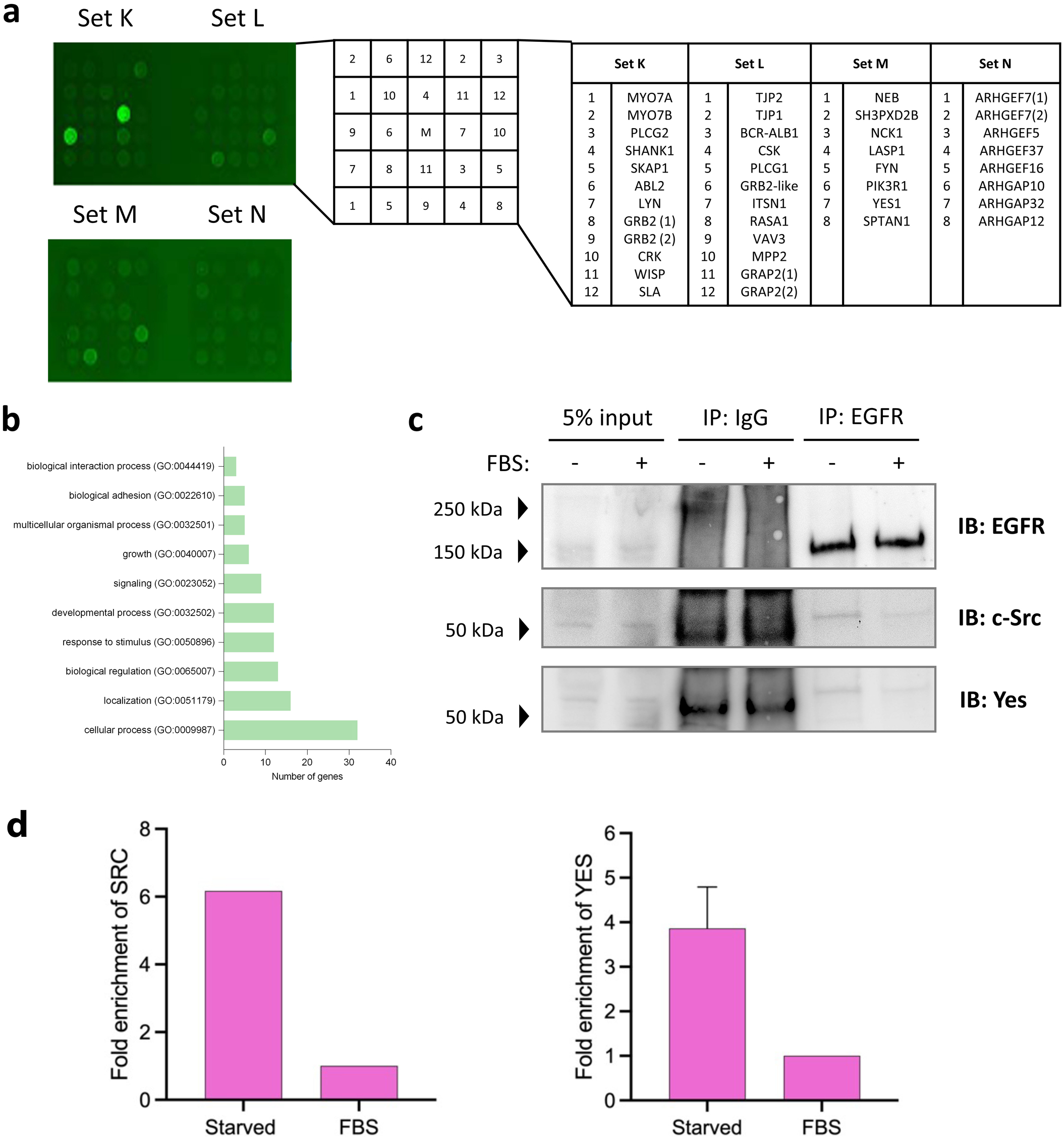
A functional overview of Src homology 3 (SH3) domain-containing PRM interactors for each of epidermal growth factor receptor (EGFR), fibroblast growth factor receptor 2 (FGFR2) and ERBB2 (HER2). **(a)** A peptide microarray demonstrating interactions between the C-terminal tail of HER2 and glutathione S-transferase (GST) fusion SH3 domain-containing proteins spotted on to a positional grid. The correlation between each protein and its grid position is shown alongside. Bound peptides are those with two fluorescent spots present. The control peptide microarray is shown in **Supp.** Fig. 1. **(b)** Overrepresented GO biological process terms amongst interacting SH3 domain-containing peptides. **(c)** Immunoprecipitation demonstrating that EGFR forms a protein complex with c-SRC and YES under serum-starved conditions (in the absence of EGFR phosphorylation **Supp.** Fig 2). Binding of c-SRC and YES is reduced in the presence of FBS. Raw data blots shown in **Supp.** Fig. 3. **(d)** Quantification of immunoprecipitated SRC (left panel) and YES (right panel): N = 3, derived from densitometry measurement of western blotting in **(c)**.

By mass spectrometry, the most commonly identified SH3 domain-containing interactor was SPTAN1 followed by multiple motility and cytoskeletal modifying myosin proteins, including modifying myosin 9 (MYH9), myosin 14 (MYH14) and myosin 6 (MYO6) proteins, as well as the unconventional myosin 18A (MY18A) protein. There were no SH3 domain-containing interactors for EGFR or HER2 in squamous lung H520 cells or for FGFR2 in the control HEK293T cells.

Amongst these proteins, the most overrepresented processes (**Fig. 4b**) related to cellular homeostasis (“cellular process (GO:0009987)”, “biological regulation (GO:0065007)”, “multicellular organismal process (GO:0032501)”, “response to stimulus (GO:0050896)”), cellular invasion and cellular migration (“localisation (GO:0051179)”, “biological adhesion (GO:0065007)”, “developmental process (GO:0032502)”). The cellular component term ‘plasma membrane’ was associated with 34 (83%) of the 41 PRM-SH3 interactors, representing statistical overrepresentation (false discovery rate, FDR 1.44x10^-10^).

The interactions reported here are fundamentally dependent on the relative concentrations of the RTKs and their binding partners and their ability to localise within a given cell to compete for the receptor PRM. Thus, environmental conditions, both outside and within the cell, will dictate the complement of binding proteins observed. Our analysis of PRM-binding ligands does not fully represent the potential for promiscuity in binding of proteins because it can only represent the relative expression levels that exist in cell lysates under the conditions of the experiment. A change in concentration of a given SH3 domain-containing protein or a RTK by a few-fold might lead to the complement of affinity purified proteins being modified. This is exemplified by the absence of binding of the previously reported binding of GRB2 and PLCƳ1 to FGFR2 in HEK293T in this study. In earlier studies the PLCƳ1-FGFR2 interaction was observed in HEK293T cells when the adaptor protein GRB2 was knocked down and the low endogenous expression of FGFR2 was enhanced by stable transfection of the receptor.^19^

### Validation of interaction between EGFR and c-SRC/YES

The SH3 domain-containing Src family proteins SRC and YES were both identified by AP-MS as EGFR interactors in HEK293T cells (**Table 2**). These proteins were selected to exemplify our interactome interactors because of extensive reported characterisation of Src family SH3 domain interactions^34^, the requirement for reduced stringency for detection, and their potential importance in being able to initiate downstream signalling (including cancer signalling) on binding to an RTK. We immunoprecipitated EGFR after an 18-hour period in which cells were cultured under serum-starved conditions without foetal bovine serum, FBS (i.e., in the absence of growth factor). These conditions are commonly reported to replicate basal, non-phosphorylated RTK conditions. The cells were then either exposed to FBS or persistently starved. The interactions of the SH3 domains from both SRC and YESwith non-phosphorylated EGFR were confirmed in serum-starved cells (**Fig. 4c**). Both SRC and YES have SH3 and SH2 domains, however serum-starvation negates receptor phosphorylation and hence SH2 binding sites (control data **Supp. Fig 2**). The addition of FBS reduces the binding of the Src family proteins to the receptor. The reason for this is not clear, however, it could reflect that SRC and YES are recruited by other receptors that are activated in the presence of low levels of stimulating ligands in FBS.

Further validation of a subset of interactions observed in Table 2 is based in the literature where interactions between the RTKs studied herein and SH3 domain-containing proteins are reported (see Biogrid (https://thebiogrid.org)). In the absence of canonical interactions between pTyr and SH2 domains these reported interactions can be inferred to be between the RTK-PRMs and SH3 domains in the respective proteins. For example, the following interactions are cited: EGFR with Myosin-9^38^; ZO-1^39^; Src substrate cortactin^40^; Sortin nexin-9^40^; SH3 domain-containing protein 19^14^; Brain-specific angiogenesis inhibitor1-associated protein 2^41^; FGFR2 with Dystonin^14^ and HER2 with Transport and Golgi organisation protein 1 homolog^14^.

## DISCUSSION

Signalling derived from RTKs is implicated in the development and progression of multiple malignancies^2^. For the most part, the oncogenic action of RTKs has classically been regarded to result from kinase activity upregulation in response to mutation or a surfeit of growth factor. Despite this, there is evidence that overexpression of wild-type RTKs also closely correlates with outcomes in a number of malignancies^2^. Whilst there is some evidence that the increased local concentration of RTKs that results from their overexpression drives signalling, the heterologous mechanisms underlying this are not well delineated. There is, nevertheless, a growing body of evidence for the presence of a diverse and functionally important, but as yet poorly mapped, RTK interactome^14^.

We have previously demonstrated that in conditions of relative RTK excess, a PRM sequence incorporated within the C-terminal tail of FGFR2 induces activation of intracellular effectors through SH3 domain-mediated interactions that occur in the absence of tyrosine kinase upregulation, and which associate with disease outcomes in a number of cancers^18–20^. We expand on this early work here by analysis of the interactome for a PRM from each of the RTKs; EGFR, FGFR2 and HER2, in cell lines derived from four cancers that together represent a diverse range of histological subtypes.

Overall, a greater number of PRM interactors were identified by affinity purification mass spectrometry for EGFR and FGFR2 than for HER2 (**Fig. 2a**). This reflects a relatively lower ability of the HER2 PRM to interact with intracellular proteins, which is supported by the small number of interactors bound by protein domain microarray (**Fig. 4a**). These data may suggest that some RTKs more freely dictate Tier 2 signalling than others. The C-terminus of HER2 includes two canonical SH3 domain binding PRMs and five other PRMs (**Supp. Table 1**) so our data presented here do not cover the entire range of possible Tier 2 interactions. Furthermore, the propensity of any given PRM interaction is likely to be dictated by differences in access to the RTK PRM that result from the conformation of the overall tail sequence within the cellular environment^42^. Underscoring this point, it should also be highlighted that, in this study we selected individual representative PRMs from the three RTKs, however each RTK does have other PRMs which could also provide binding sites (see **Supp. Table 1**). Therefore, further analysis of other PRMs might reveal a larger role for HER2 in Tier 2 signalling.

The interaction of a given domain with a PRM in cells is entirely dependent on its relative concentration with respect to other competing domains and its cellular localisation. Interestingly, our data does appear to show a connection between the intracellular concentration of domain-containing proteins that dictates the likelihood of their PRM-mediated interaction. This is in keeping with our previous work and is coupled with evidence provided here that a majority of interacting these proteins are found at the cell surface. This highlights a complex signalling dynamic in which both the specific RTK PRM and the local cellular environment in which it sits direct signalling outcomes.

Amongst the interactors identified by mass spectrometry for EGFR, FGFR2 and HER2, contributors to metabolic, homeostatic and migratory processes were overrepresented. Whilst this was broadly reflected by the specific pathways overrepresented amongst the interactor peptides, additional immune processes were seen for at least the EGFR C-terminal PRM in U251 GBM cells. The propensity of these processes resulting might suggest that in non-pathological conditions Tier 2 signalling is responsible for ‘house-keeping’ and ‘response to environmental stress’ functions which can be fine-tuned and potentially reversed. This is in contrast to Tier 1 signals that tend to result in profound and irreversible cellular outcomes such as differentiation, proliferation and cell death.

Given that we have previously demonstrated a specific role for the SH3 domain in mediating interactions with RTK C-terminal PRMs, we sought to additionally identify interactors with this domain for each of our studied RTKs. In doing so, we characterised 41 SH3 domain-containing proteins of potential interest. Amongst these, processes relating to migration, invasion and homeostasis were overrepresented. More broadly, the overrepresentation of migratory and homeostatic terms amongst the identified interacting proteins may provide evidence for the ability of cancer to hijack an evolutionarily conserved mechanism. It would, for example, be beneficial for cells lacking in growth factor stimulation to upregulate survival pathways and potentially even migrate to an area of greater RTK ligand availability or escape unfavourable environmental conditions. This would occur via the Tier 2 mechanism demonstrated here but could potentially also be hijacked by a cancer cell through upregulation of RTK and/or downstream effector expression in response to a stressor that results in a relative excess of RTK tail PRM regions. Indeed, it is possible that the change in expression profile of possible Tier 2 signal-initiating effectors could be stimulated by cellular response to therapeutic intervention, hence providing a mechanism for resistance.

In support of the presence of this mechanism, many of the specific interactors to the studied RTKs are already recognised to contribute to adverse cancer outcomes. We have, for instance, validated an interaction between the EGFR PRM and the SH3-proteins SRC and YES. Dysregulation of the Src family kinases is well recognised in cancer and contributes to poorer outcomes^24^. This has typically been regarded to result from increased EGFR transactivation, but the work here suggests an additional mechanism through which the EGFR/Src family kinase proteins may contribute to deleterious cancer outcomes. Favouring signalling via this mechanism, lipid rafts have been shown to provide a platform for EGFR and c-SRC interaction in breast cancer cells^36^.

Another interactor of potential interest is the EVH1 domain-containing switch associated protein 70 (SWAP70), which associated with EGFR in GBM U251 cells. This has been shown to mediate GBM migration and invasion by regulating CD44 expression^43^. Likewise, the SH3 domain-containing protein MYCBP2 is linked to mitotic fate and thereby chemoresistance in an HEK293T cell population that is similar to the line that MYCPB2 was pulled down from here.

This study provides evidence for the basis of Tier 2 interactions which, along with the limited number of these interactions that have now been comprehensively evaluated, require further validation. Furthermore, the weak and transient nature of SH3-PRM interactions is such that only a limited number of SH3 domain-containing proteins could be identified with high levels of confidence given the reliance of an AP/MS approach on stable and reasonably strong interactions. To this end, it would be appropriate to build on the work shown here with additional functional studies and by correlating the expression of relevant SH3 domain-containing proteins with survival in existing clinical datasets.

Our data, therefore, add considerably to contemporary developments in our understanding of the physical and functional interactome of human RTKs. Furthermore, we highlight that SH3 domains are likely to play a far more important and independent role in intracellular signalling than generally considered. In the absence of the requirement for an on/off functionality represented by tyrosine phosphorylation in SH2 domain-mediated signalling, signalling is dependent on concentration fluctuations of SH3 domain-containing proteins. This suggests roles in responding to environmental stress and metabolic and homeostatic regulation. However, under conditions of aberrant or prolonged response, oncogenic signalling can prevail. In the light of this, we might need to consider alternative therapeutic approaches to cancer and directed kinase inhibitor resistance.

The weak and transient nature of SH3-PRM interactions means that they are hard to identify using previously adopted protocols and stringency ‘cut-offs’ for proteomics studies. Therefore, this study is likely to underrepresent the breadth and number of interactions maintained by the RTK PRM. There are also clear but previously under-recognised differences in the ability of AP-MS scoring systems to identify these interactions, with a greater number of possible interactors identified in this study using Perseus rather than SAINTexpress. Clearly, this study is limited to PRMs from a subset of RTKs and five cell lines under one set of conditions. To truly reflect the possible interactions of the RTK-PRM transcriptome a more substantial screen would be required which would include multiple cell lines, representative physiological conditions and extensive validation of physiological and pathological relevance.

## METHODS

### Materials and reagents

RIPA Lysis and Extraction Buffer (#89900) was obtained from Thermo Fisher Scientific. Bovine serum albumin (BSA; A9647) was purchased from Merck.

### Mammalian cell culture

Human cell lines representing BrAC (SK-BR-3, ATCC HTB-30™) and LSCC (NCI-H520, ATCC HTB-182™), in addition to a control human embryonic kidney (HEK)-293T cell line, were obtained from the American Type Culture Collection (ATCC; Virginia, USA). The highly-transfectable HEK293T cell line was generated by stably transfecting HEK293T cells with FGFR2, as has previously been described^17^. This cell line has been extensively utilised to study the impact of the upregulation of FGFR2 and other endogenously expressed RTKs^44^. Human cell lines representing OAC (OE19, JROECL19) and GBM (U251 MG, #89081403) were obtained from the European Collection of Cell Cultures (ECACC; Salisbury, UK). All cells were assessed for mycoplasma contamination at monthly intervals using the LookOut Mycoplasma Detection kit (MP0035, Thermo Fisher Scientific, Loughborough, UK).

Each cell line was maintained as a sub-confluent culture at 37°C in a humidified atmosphere with 5% carbon dioxide (CO2) in air. OE19 and H520 cells were maintained in Roswell Park Memorial Institute (RPMI)-1640 growth medium (R6504, Sigma Aldrich, St Louis., USA) supplemented with 10% (v/v) foetal bovine serum (FBS; Sigma Aldrich) and 2mM L-glutamine. SK-BR-3, U251-MG and HEK293T cells were cultured in Dulbecco’s Modified Eagle Medium (DMEM) supplemented with 10% (v/v) FBS, 50μg/ml gentamicin and 7μg/ml puromycin (Sigma Aldrich). When not passaged, 50% media exchanges were undertaken at three-day intervals.

### Cell lysate and sample preparation

Prior to lysis and streptavidin pulldown using wild-type or scrambled RTK C-terminal tail sequences, cells were grown to around 90% confluence in a 100 mm tissue culture dish. In order to maintain cellular viability, each cell line was maintained in RPMI/DMEM supplemented with 10% FBS, as outlined above. Though FBS does contain growth factors and may therefore facilitate classic ligand inducible RTK activation, RTK loop phosphorylation in the presence of FBS is nevertheless known to be low and conditions of serum (i.e., FBS) starvation are associated with cellular stress (e.g., FGFR2^15^). Furthermore, none of the wild-type or scrambled bait peptides contained ligand-inducible tyrosine residues, thereby favouring PRM-mediated interactions even in the presence of supplemented growth factor.

Prior to lysis, cells were washed three times in ice-cold phosphate buffered saline (PBS) and harvested in RIPA Lysis & Extraction buffer comprising 1% NP40 (GenTex, Irvine), 1% Na-Deoxycholate (Sigma-Aldrich, St. Louis), 0.1% SDS (Sigma-Aldrich, St. Louis), 0.15M NaCl (Honeywell, Seelze), 0.01M Na-phosphate (Sigma-Aldrich, St. Louis), 2mM EDTA (Sigma-Aldrich, St. Louis), 50mM NaF (Sigma-Aldrich, St. Louis) and 0.2mM Na-orthovanadate (Sigma-Aldrich, St. Louis) at pH 7.2. Cells were further disrupted and homogenised via hydrodynamic shearing using a 0.8mm needle followed by one hour of continuous rotation at 4°C. Cell debris and DNA were subsequently removed through aspiration of the supernatant following by centrifugation at 2000g for 15 minutes at 4°C. Sixteen cell pellets were independently prepared for each cell line. Protein concentration was determined using a colorimetric Pierce™ BSA Protein Assay Kit (#23225, Thermo Fisher Scientific). Lysates were subsequently stored at -80°C prior to experimentation.

### Biotinylated bait peptides

The presence of large affinity tags, particularly when fused to small peptide sequences, can impact on the structure and therefore function of the proteins to which they are bound^45^. In contrast, conjugation of biotin is unlikely to impact on function given its small size. Synthetic peptides representing the PRM-containing cytoplasmic tail sequence of EGFR (VQNPVFHNQPLNPAPSRDPH – residues 1105-1124), HER2 (DVRPQPPSPREGPLPAAR – residues 1144-1161) and FGFR2 (EPSLPQFPHINGSVKT – residues 806-821) were therefore commercially prepared and modified through the covalent N-terminal addition of a biotin tag (GenScript Biotech, The Netherlands). Scrambled control sequences that did not contain a PRM were similarly prepared for EGFR (VQNLVFHNQLLNLALSRDLH), HER2 (DVRLQLLSLREGLLLAAR) and FGFR2 (ELSLLQFLHINGSVKT). Tyrosine residues (Y1110: EGFR and Y812: FGFR2) in RTK sequences were mutated to phenyl alanine (F1110 and Y812 respectively) to remove any opportunity for phosphorylation. In all cases, peptide sequences were separated from the biotin tag by two inert, highly hydrophilic polyethylene glycol (PEG) spacers in order to increase solubility. PEG spacers are known to have minimal impact on the conformational properties of small neutral peptides^46^.

### Immunoprecipitation and affinity enrichment

Each biotinylated tail sequence and matched scrambled control was assayed in biological triplicate in each cell line. Protein extracts were pre-cleared through the addition of 1mg total protein lysate to 10µl streptavidin agarose (Pierce, 88817) beads for a period of one hour. Streptavidin beads for protein elution were pre-incubated with 50µg peptide in 100µl RIPA buffer at 4°C for one hour. The streptavidin beads were removed from the pre-cleared protein solution by centrifugation and the pre-incubated streptavidin beads subsequently added. Samples were incubated overnight at 4°C with constant agitation. Following this, the streptavidin beads were pelleted by centrifugation and the supernatant discarded. The now protein-bound beads were subsequently washed twice in RIPA buffer and the beads stored at -80°C.

Protein elution proceeded though the incubation of streptavidin beads with 30μl 20mM dithiothreitol in a 5% sodium dodecyl sulphate (SDS), t0mM Tris-HCl (pH 7.6) buffer at 90°C for 10 minutes. Proteins were alkylated through the subsequent incubation for 30 minutes in the dark with iodoacetamide to a final concentration of 150mM. Samples were then prepared for mass spectrometry by protein tryptic digest using the Suspension Trapping (Strap) method for bottom-up proteomics analysis, as has previously been described^45^.

### Liquid chromatography mass spectrometry

Processed peptides were analysed by nanoflow liquid chromatography mass spectrometry (LC-MS) using an EASY-nLC 1000 Liquid Chromatograph (Thermo Fisher Scientific) connected to a custom-made 30-cm capillary emitter column (75μm inner diameter, 3μm Reprosil-Pur 120 C18 media). Mass spectrometry analysis was performed on a linear quadrupole ion trap - orbitrap (LTQ-Orbitrap) Velos mass spectrometer (Thermo). Total acquisition time was set to 100 minutes, with a gradient of 3-22% acetonitrile in 0.1% formic acid. For the survey scan, the resolving power was set at 60,000 with a scan range of 305-1350 amu. MS/MS data were obtained by fragmenting up to the twenty most intense ions in the linear ion trap. Data were searched against the Uniprot human protein sequence database with MaxQuant software package (www.maxquant.org)^47^. The maximum protein and peptide false discovery rates were set to 0.01.

### Probabilistic modelling for scoring AP/MS data

Significance Analysis of INTeractome express (SAINTexpress) software was used to estimate the probability that each postulated bait-prey protein-protein interaction from the AP/MS data was true.^48^ This probabilistic model is not vulnerable to quantitative variation of prey proteins across studied purifications and additionally accounts for negative control purifications such as those used here; thereby robustly removing background noise whilst accommodating for the impact of random sampling. A final interaction score (AvgP) for each bait-prey protein-protein pair is then calculated by averaging the probabilities for individual replicates.

Here, AP-MS data for each bait peptide were examined separately using SAINTexpress version 3.1.0 (http://saint-apms.sourceforge.net). Final AvgP results of 0.5 or greater were retained for further analysis. Common contaminants were removed using the peer-annotated Contaminant Repository for Affinity Purification-Mass Spectrometry data (CRAPome), version 2.0 (http://crapome.org)^49^. This uses mass spectrometry data from 716 experiments to filter possible contaminants. Only proteins with a CRAPome frequency of less than 358/716 (50%) were retained. These interactors were subsequently explored using functional annotation. A separate group of low confidence interactors (LCIs) with an AvgP score of greater than 0 but less than 0.5 and a CRAPome frequency of less than 50% were also identified for analyses relating to the interaction of SH3 domain-containing proteins with PRMs.

A second approach to the identification of SH3-containing binding partners was undertaken using Perseus (2.0.3.0)^37^. This computational platform uses peptide intensity-based quantification to identify proteins that are enriched in the presence of specific bait peptides, with a permutation-based false discovery rate applied for each sample-control pair. Interactors were identified here using a two-sample t-test with an FDR cut-off of 0.05.

### Functional annotation

Protein domains were manually annotated using HumanMine v12^50^. Conserved Pfam protein domains present in each HCI were identified using the Ensembl BioMart data mining tool (https://uswest.ensembl.org/info/data/biomart/index.html)^51^. Identified Pfam domains were grouped into Clans^52^. Searches were restricted to superfamilies. Over-represented Gene Ontology (GO) terms were identified from interactors using Protein Analysis Through Evolutionary Relationships (PANTHER) version 16.0^53^. Functionally enriched GO Biological Process (BP), Molecular Function (MF) and Cellular Component (CC) terms were identified using a reference *Homo Sapiens* gene set.

### Protein domain microarray

A protein-domain microarray was used in order to identify potential SH3 domain-containing interactors with the HER2 receptor. The use of this system to identify novel protein-protein interactions has been described previously^54^. Briefly, the experimental workflow includes purification of glutathione S-transferase (GST) fusion SH3 domain-containing proteins, the arraying of these proteins on a microarray and their probing using PRM-containing RTK tail sequences with interactions determined using fluorescent probes.

### Purification of GST fusion proteins

Overexpression of glutathione S-transferase (GST) fusion proteins was induced in DH5α *Escherichia coli* cells (Life Technologies, MD, USA) using 0.4mM isopropyl β-D-thiogalactopyranoside and cells subsequently broken by sonication. GST fusion proteins were extracted from the resultant lysates by centrifugation at 12000 *g* for 10 minutes followed by binding to glutathione-Sepharose 4B beads (Amersham Pharmacia Biotech, NJ, USA). Purified proteins were eluted using 30mM glutathione, 50mM Tris/HCl, pH 7.5 and 120mM NaCl then stored at -70°C.

### Protein microarray peptides

A protein microarray incorporating 30 SH3 domain-containing protein sequences was generated as outlined previously.^55^ Approximately 250ng of each protein stock was arrayed on to one of 25 specific spots on a nitrocellulose pre-coated glass FAST™ slide (Schleicher & Schuell, NH, USA). Spots were spaced at a 700µm distance from one another, and each protein was spotted in duplicate then allowed to air dry. A control GST-alone spot was placed in the centre of this grid.

### Protein microarray probes

Biotinylated peptides representing the PRM-containing HER2 tail sequence (GGGGAAPQPHPPPAFSPAFDNL) and the non-PRM IGF1R (GGGGRKNERALPLPQSST) tail sequence were synthesised by Genscript (NJ, USA). Each was then bound to 5µl Cy3-streptavidin (Fluorolink™; Amersham Pharmacia Biotech) in 500µl PBS containing 0.1% Tween 20; PBST) and then incubated with 20µl biotin-agarose beads (Sigma, MO, USA).

### Probe-peptide interaction

Arrayed slides were blocked in PBST containing 3% (w/v) powdered milk within an Atlas Glass Hybridisation Chamber (Clontech, CA, USA) then hybridised to 400µl fluorophore-tagged peptide for 1 hour. Three 10 minute washes with PBST were subsequently used to remove unbound peptide and the slide dried via centrifugation. Following this, a 550nm long pass filter was used for the detection of the Cy3-labelled probes via a GeneTAC™ LSIV scanner (Genomic Solutions). Flourescence of two dots representing the same protein is regarded as a positive indication of peptide-probe interaction.

### Immunoprecipitation and western blotting

HEK 293T cells were grown to 80% confluency and then cultured in media not supplemented with FBS for 18 hours. Following this period, cells were cultured for 45 minutes with and without FBS supplementation. Cells were lysed in lysis buffer (50mM HEPES, 50mM NaCl 1mM EGTA, 10% (w/v) glycerol, 1mM sodium orthovanadate, 10mM sodium fluoride, 0.1% NP-40, supplemented with protease inhibitors) and cleared of cell debris via centrifugation.

Following quantification of protein concentration, 1mg of cell lysate was incubated at room temperature with 10µg anti-EGFR (SCBT; sc-120-AC) or 10µg anti-mouse IgG (SCBT; sc-2343) for 2 hours at room temperature, with gentle rotation. Immunoprecipitants were subsequently washed three times with 1ml lysis buffer. Samples were then analysed by western blotting. Antibodies used in western blotting were EGFR (CST; cat no. 4267), Src (CST; cat no. 2123), Yes (CST; cat no. 3201).

### Correlation of binding patterns with expression profiles

In order to determine whether the interactors identified via AP-MS correlated with the specific expression profile of these interactors in each cell line, we extracted mRNA expression data from the Cancer Cell Line Encyclopedia (CCLE)^56,57^. Specifically, gene expression transcript per million (TPM) values of protein coding genes for all DepMap cell lines, reported using a pseudo-count of log2(TPM+1), were downloaded from DepMap Public 23Q2 primary files (file: ‘OmicsExpressionProteinCodingGenesTPMLogp1’). A detailed description of the pipelines used to generate these expression data can be found at https://github.com/broadinstitute/ccle_processing#rnaseq. Data relating to the studied cell lines were extracted using the following DepMapIDs: ACH-000017 (SK-BR-3), ACH-000679 (OE19), ACH-000049 (HEK293T), ACH-000232 (U251) and ACH-000395 (NCIH520). We then searched within these datasets for genes corresponding to all SH3-domain proteins identified to be a possible interactor in at least one cell line (these are summarised in **Table 2**). The pseudo-count of these genes is shown for each cell line and correlated against the presence or absence of an interaction with any of the studied RTKs in that cell line (i.e. counts are provided either for ‘no interaction’, in which the specific interactor from the studied cohort did not bind to any of the RTKs studied in that cell line or ‘interaction’, in which the specific interactor bound to at least one of the RTKs in that cell line.

### Illustrations

Illustrations were created using Servier Medical Art, provided by Servier, licensed under a Creative Attribution 3.0 unreported license. Unless otherwise stated, graphs have been generated using GraphPad Prism 8.1.2. (GraphPad Software, CA, USA).

## ACKNOWLEDGEMENTS

The authors are grateful to Prof. Niamh Forde and Dr Chiara Francavilla for their feedback on earlier versions of this manuscript. The authors are in addition grateful to Dr. Cari Sagum for their assistance in the protein domain array experiments.

## AUTHOR CONTRIBUTIONS

JEL devised the study and led experimental design. CMJ and AR contributed to study design. AR, CMJ, KMS, SZM and SK collected data. JEL, AR, CMJ, and SK contributed to the analysis and interpretation of data. MTB designed and performed the SH3 protein domain array experiments. CMJ performed all bioinformatic analyses. CMJ authored the first draft of the manuscript. All authors contributed to subsequent revisions of the draft manuscript.

## ETHICS DECLARATION

Review and/or approval by an ethics committee was not needed for this study because no experiments or data acquisition involved patients or patient samples.

## DATA AVAILABILITY

Mass spectrometry proteomics data have been deposited to the ProteomeXchange Consortium via the Proteomics IDEntifications Database (PRIDE) partner data repository with the dataset identifier PXD041383.

## REFERENCES

1. Lemmon, M. A. & Schlessinger, J. Cell signaling by receptor tyrosine kinases. Cell 141, 1117–1134 (2010).

2. Du, Z. & Lovly, C. M. Mechanisms of receptor tyrosine kinase activation in cancer. Mol Cancer 17, 58 (2018).

3. Bhullar, K. S. et al. Kinase-targeted cancer therapies: progress, challenges and future directions. Molecular Cancer 17, 48 (2018).

4. Tong, A. H. Y. et al. A combined experimental and computational strategy to define protein interaction networks for peptide recognition modules. Science 295, 321–324 (2002).

5. Kazlauskas, A. et al. Large-scale screening of preferred interactions of human Src homology-3 (SH3) domains using native target proteins as affinity ligands. Mol Cell Proteomics 15, 3270– 3281 (2016).

6. Xin, X. et al. SH3 interactome conserves general function over specific form. Mol Syst Biol 9, 652 (2013).

7. Liu, B. A., Engelmann, B. W. & Nash, P. D. High-throughput analysis of peptide-binding modules. Proteomics 12, 1527–1546 (2012).

8. Carducci, M. et al. The protein interaction network mediated by human SH3 domains. Biotechnol Adv 30, 4–15 (2012).

9. Wu, C. et al. Systematic identification of SH3 domain-mediated human protein-protein interactions by peptide array target screening. Proteomics 7, 1775–1785 (2007).

10. Landgraf, C. et al. Protein interaction networks by proteome peptide scanning. PLoS Biol 2, E14 (2004).

11. Zarrinpar, A., Bhattacharyya, R. P. & Lim, W. A. The structure and function of proline recognition domains. Sci STKE 2003, RE8 (2003).

12. Kay, B. K., Williamson, M. P. & Sudol, M. The importance of being proline: the interaction of proline-rich motifs in signaling proteins with their cognate domains. FASEB J 14, 231–241 (2000).

13. Brannetti, B., Via, A., Cestra, G., Cesareni, G. & Helmer-Citterich, M. SH3-SPOT: an algorithm to predict preferred ligands to different members of the SH3 gene family. J Mol Biol 298, 313–328 (2000).

14. Salokas, K. et al. Physical and functional interactome atlas of human receptor tyrosine kinases. EMBO Rep e54041 (2022) doi:10.15252/embr.202154041.

15. Teyra, J. et al. Comprehensive analysis of the human SH3 domain family reveals a wide variety of non-canonical specificities. Structure 25, 1598–1610.e3 (2017).

16. Kurochkina, N. & Guha, U. SH3 domains: modules of protein-protein interactions. Biophys Rev 5, 29–39 (2013).

17. Lin, C.-C. et al. Inhibition of basal FGF receptor signaling by dimeric Grb2. Cell 149, 1514–1524 (2012).

18. Ahmed, Z. et al. Grb2 monomer-dimer equilibrium determines normal versus oncogenic function. Nat Commun 6, 7354 (2015).

19. Timsah, Z. et al. Competition between Grb2 and Plcγ1 for FGFR2 regulates basal phospholipase activity and invasion. Nat Struct Mol Biol 21, 180–188 (2014).

20. Timsah, Z. et al. Grb2 depletion under non-stimulated conditions inhibits PTEN, promotes Akt-induced tumor formation and contributes to poor prognosis in ovarian cancer. Oncogene 35, 2186–2196 (2016).

21. Bornet, O. et al. Identification of a Src kinase SH3 binding site in the C-terminal domain of the human ErbB2 receptor tyrosine kinase. FEBS Lett 588, 2031–2036 (2014).

22. Ball, L. J., Kühne, R., Schneider-Mergener, J. & Oschkinat, H. Recognition of proline-rich motifs by protein-protein-interaction domains. Angew Chem Int Ed Engl 44, 2852–2869 (2005).

23. Lopez-Gines, C. et al. New pattern of EGFR amplification in glioblastoma and the relationship of gene copy number with gene expression profile. Mod Pathol 23, 856–865 (2010).

24. Selvaggi, G. et al. Epidermal growth factor receptor overexpression correlates with a poor prognosis in completely resected non-small-cell lung cancer. Ann Oncol 15, 28–32 (2004).

25. Hirsch, F. R., Varella-Garcia, M. & Cappuzzo, F. Predictive value of EGFR and HER2 overexpression in advanced non-small-cell lung cancer. Oncogene 28 **Suppl 1**, S32–37 (2009).

26. Ménard, S. et al. HER2 overexpression in various tumor types, focussing on its relationship to the development of invasive breast cancer. Ann Oncol 12 **Suppl 1**, S15–19 (2001).

27. Masuda, H. et al. Role of epidermal growth factor receptor in breast cancer. Breast Cancer Res Treat 136, 331–345 (2012).

28. Koka, V. et al. Role of Her-2/neu overexpression and clinical determinants of early mortality in glioblastoma multiforme. Am J Clin Oncol 26, 332–335 (2003).

29. Hu, Y. et al. HER2 amplification, overexpression and score criteria in esophageal adenocarcinoma. Mod Pathol 24, 899–907 (2011).

30. Klempner, S. J. et al. FGFR2-Altered Gastroesophageal Adenocarcinomas Are an Uncommon Clinicopathologic Entity with a Distinct Genomic Landscape. Oncologist 24, 1462–1468 (2019).

31. Santolla, M. F. & Maggiolini, M. The FGF/FGFR System in Breast Cancer: Oncogenic Features and Therapeutic Perspectives. Cancers (Basel) 12, 3029 (2020).

32. Jimenez-Pascual, A. & Siebzehnrubl, F. A. Fibroblast Growth Factor Receptor Functions in Glioblastoma. Cells 8, 715 (2019).

33. Barretina, J. et al. The Cancer Cell Line Encyclopedia enables predictive modelling of anticancer drug sensitivity. Nature 483, 603–607 (2012).

34. Ladbury, J. E. & Arold, S. Searching for specificity in SH domains. Chem Biol 7, R3–8 (2000).

35. Belli, S. et al. c-Src and EGFR inhibition in molecular cancer therapy: What else can we improve? Cancers (Basel) 12, 1489 (2020).

36. Irwin, M. E., Bohin, N. & Boerner, J. L. Src family kinases mediate epidermal growth factor receptor signaling from lipid rafts in breast cancer cells. Cancer Biol Ther 12, 718–726 (2011).

37. Tyanova S et al. The Perseus computational platform for comprehensive analysis of (prote)omics data. Nat Methods 13, 731–740 (2016)

38. Chiu, H. C. et al. EGFR and myosin II inhibitors cooperate to suppress EGFR-T790M-mutant NSCLC cells. Mol. Oncol. 6, 299–310 (2012)

39. Kaihara, T. et al. Redifferential and ZO-1 reexpression in liver-metastasized colorectal cancer: Possible association with epidermal growth factor receptor-induced tyrosine phosporylation of ZO-1. Cancer Sci. 94, 166–172 (2003).

40. Petschnigg, J. et al. The mammalian-membrane two-hybrid assay (MaMTH) for probing membrane-protein interactions in human cells Nature Meth. 11, 585–592 (2014).

41. Li, J. et al. Perturbation of the mutated EGFR interactiome identifies vulnerabilities and resistance mechanisms *Mol*. Syst. Biol. 9, 705 (2013)

42. Pinet, L. et al. Structural and dynamic characterization of the C-terminal tail of ErbB2: Disordered but not random. Biophys. J. 120, 1869–1882 (2021).

43. Shi, L. et al. SWAP-70 promotes glioblastoma cellular migration and invasion by regulating the expression of CD44s. Cancer Cell Int. 19, 305 (2019).

44. Ahmed, Z., Schüller, A. C., Suhling, K., Tregidgo, C. & Ladbury, J. E. Extracellular point mutations in FGFR2 elicit unexpected changes in intracellular signalling. Biochem J 413, 37–49 (2008).

45. Zougman, A., Selby, P. J. & Banks, R. E. Suspension trapping (STrap) sample preparation method for bottom-up proteomics analysis. Proteomics 14, 1006–1000 (2014).

46. Xue, Y., O’Mara, M. L., Surawski, P. P. T., Trau, M. & Mark, A. E. Effect of poly(ethylene glycol) (PEG) spacers on the conformational properties of small peptides: a molecular dynamics study. Langmuir 27, 296–303 (2011).

47. Cox, J. & Mann, M. MaxQuant enables high peptide identification rates, individualized p.p.b.-range mass accuracies and proteome-wide protein quantification. Nat. Biotechnol. 26, 1367– 1372 (2008).

48. Teo, G. et al. SAINTexpress: improvements and additional features in Significance Analysis of INTeractome software. J. Proteomics 100, 37–43 (2014).

49. Mellacheruvu, D. et al. The CRAPome: a contaminant repository for affinity purification-mass spectrometry data. Nat. Methods 10, 730–736 (2013).

50. Smith, R. N. et al. InterMine: a flexible data warehouse system for the integration and analysis of heterogeneous biological data. Bioinformatics 28, 3163–3165 (2012).

51. Smedley, D. et al. BioMart--biological queries made easy. BMC Genomics 10, 22 (2009).

52. El-Gebali, S. et al. The Pfam protein families database in 2019. Nucleic Acids Res. 47, D427–D432 (2019).

53. Mi, H., Muruganujan, A. & Thomas, P. D. PANTHER in 2013: modeling the evolution of gene function, and other gene attributes, in the context of phylogenetic trees. Nucleic Acids Res. 41, D377–386 (2013).

54. Chen, J., et al. Protein domain microarrays as a platform to decipher signaling pathways and the histone code. Methods 184, 4–12 (2020).

55. Yang, Y. et al. TDRD3 is an effector molecule for arginine-methylated histone marks. Mol. Cell 40, 1016–23 (2010).

56. Barretina, J, et al. The Cancer Cell Line Encyclopedia enables predictive modelling of anticancer drug sensitivity. Nature 483, 603–307 (2012).

57. Ghandi, M., et al. Next-generation characterization of the Cancer Cell Line Encyclopedia. Nature 569, 503–508 (2019).

