## Supplemental Table 1 for "Affinity purification mass spectrometry characterization of the interactome of receptor tyrosine kinase proline rich motifs in cancer"

**Supplementary Table 1: Proline-rich motifs and other, less common, Src homology 3 (SH3) domain-binding sequences within the cytoplasmic tails of all 58 receptor tyrosine kinases (RTKs) encoded by the human genome.** Common SH3 domain-binding RTK tail sequences (PxxP) are highlighted in yellow. Cannonical PxxPxR, RxPxxP and RxxPxxP motifs are underlined. Less common SH3 domain-binding RTK tail sequences are shown in purple (PxxDY), blue (PxPxxR) and green (R/KxLP). x can be any amino acid.

| **Type 1 RTKs (epidermal growth factor receptor family)** | |
| --- | --- |
| EGFR: epidermal growth factor receptor | VIQGDERMHLPSPTDSNFYRALMDEEDMDDVVDADEYLIPQQGFFSSPSTSRTPLLSSLSATSNNSTVACIDRNGLQSCPIKEDSFLQRYSSDPTGALTEDSIDDTFLPVPEYINQSVPKRPAGSVQNPVYHNQPLNPAPSRDPHYQDPHSTAVGNPEYLNTVQPTCVNSTFDSPAHWAQKGSHQISLDNPDYQQDFFPKEAKPNGIFKGSTAENAEYLRVAPQSSEFIGA |
| HER2: erb-b2 RTK 2 | VIQNEDLGPASPLDSTFYRSLLEDDDMGDLVDAEEYLVPQQGFFCPDPAPGAGGMVHHRHRSSSTRSGGGDLTLGLEPSEEEAPRSPLAPSEGAGSDVFDGDLGMGAAKGLQSLPTHDPSPLQRYSEDPTVPLPSETDGYVAPLTCSPQPEYVNQPDVRPQPPSPREGPLPAARPAGATLERPKTLSPGKNGVVKDVFAFGGAVENPEYLTPQGGAAPQPHPPPAFSPAFDNLYYWDQDPPERGAPPSTFKGTPTAENPEYLGLDVPV |
| HER3: erb-b2 RTK 3 | RMARDPPRYLVIKRESGPGIAPGPEPHGLTNKKLEEVELEPELDLDLDLEAEEDNLATTTLGSALSLPVGTLNRPRGSQSLLSPSSGYMPMNQGNLGESCQESAVSGSSERCPRPVSLHPMPRGCLASESSEGHVTGSEAELQEKVSMCRSRSRSRSPRPRGDSAYHSQRHSLLTPVTPLSPPGLEEEDVNGYVMPDTHLKGTPSSREGTLSSVGLSSVLGTEEEDEDEEYEYMNRRRRHSPPHPPRPSSLEELGYEYMDVGSDLSASLGSTQSCPLHPVPIMPTAGTTPDEDYEYMNRQRDGGGPGGDYAAMGACPASEQGYEEMRAFQGPGHQAPHVHYARLKTLRSLEATDSAFDNPDYWHSRLFPKANAQRT |
| HER4: erb-b2 RTK4 | VIQGDDRMKLPSPNDSKFFQNLLDEEDLEDMMDAEEYLVPQAFNIPPPIYTSRARIDSNRSEIGHSPPPAYTPMSGNQFVYRDGGFAAEQGVSVPYRAPTSTIPEAPVAQGATAEIFDDSCCNGTLRKPVAPHVQEDSSTQRYSADPTVFAPERSPRGELDEEGYMTPMRDKPKQEYLNPVEENPFVSRRKNGDLQALDNPEYHNASNGPPKAEDEYVNEPLYLNTFANTLGKAEYLKNNILSMPEKAKKAFDNPDYWNHSLPPRSTLQHPDYLQEYSTKYFYKQNGRIRPIVAENPEYLSEFSLKPGTVLPPPPYRHRNTVV |
| **Type 2 RTKs (insulin receptor family)** | |
| InsR: insulin receptor | PEVSFFHSEENKAPESEELEMEFEDMENVPLDRSSHCQREEAGGRDGGSSLGFKRSYEEHIPYTHMNGGKKNGRILTLPRSNPS |
| IGF1R: insulin-like growth factor 1 receptor | YYSEENKLPEPEELDLEPENMESVPLDPSASSSSLPLPDRHSGHKAENGPGPGVLVLRASFDERQPYAHMNGGRKNERALPLPQSSTC |
| IRR: insulin-related receptor | RLLSFYYSPECRGARGSLPTTDAEPDSSPTPRDCSPQNGGPGH |
| **Type 3 RTKs (PDGFR, CSFR, Kit, FLT3 receptor family)** | |
| PDGFRα: platelet-derived growth factor receptor α | PGQYKKSYEKIHLDFLKSDHPAVARMRVDSDNAYIGVTYKNEEDKLKDWEGGLDEQRLSADSGYIIPLPDIDPVPEEEDLGKRNRHSSQTSEESAIETGSSSSTFIKREDETIEDIDMMDDIGIDSSDLVEDSFL |
| PDGFRβ: platelet-derived growth factor receptor β | GEGYKKKYQQVDEEFLRSDHPAILRSQARLPGFHGLRSPLDTSSVLYTAVQPNEGDNDYIIPLPDPKPEVADEGPLEGSPSLASSTLNEVNTSSTISCDSPLEPQDEPEPEPQLELQVEPEPELEQLPDSGCPAPRAEAEDSFL |
| Kit: KIT proto-oncogene RTK | STNHIYSNLANCSPNRQKPVVDHSVRINSVGSTASSSQPLLVHDDV |
| CSFR: colony-stimulating factor 1 receptor | QEQAQEDRRERDYTNLPSSSRSGGSGSSSSELEEESSSEHLTCCEQGDIAQPLLQPNNYQFC |
| FLT3: fms related tyrosine kinase 3 | GCQLADAEEAMYQNVDGRVSECPHTYQNRRPFSREMDLGLLSPQAQVEDS |
| **Type 4 RTKs (vascular endothelial growth factor receptor) family** | |
| VEGFR-1: fms related tyrosine kinase 1 | QANVQQDGKDYIPINAILTGNSGFTYSTPAFSEDFFKESISAPKFNSGSSDDVRYVNAFKFMSLERIKTFEELLPNATSMFDDYQGDSSTLLASPMLKRFTWTDSKPKASLKIDLRVTSKSKESGLSDVSRPSFCHSSCGHVSEGKRRFTYDHAELERKIACCSPPPDYNSVVLYSTPPI |
| VEGFR-2: kinase insert domain receptor | LLQANAQQDGKDYIVLPISETLSMEEDSGLSLPTSPVSCMEEEEVCDPKFHYDNTAGISQYLQNSKRKSRPVSVKTFEDIPLEEPEVKVIPDDNQTDSGMVLASEELKTLEDRTKLSPSFGGMVPSKSRESVASEGSNQTSGYQSGYHSDDTDTTVYSSEEAELLKLIEIGVQTGSTAQILQPDSGTTLSSPPV |
| VEGFR-3: fms related tyrosine kinase 4 | QGRGLQEEEEVCMAPRSSQSSEEGSFSQVSTMALHIAQADAEDSPPSLQRHSLAARYYNWVSFPGCLARGAETRGSSRMKTFEEFPMTPTTYKGSVDNQTDSGMVLASEEFEQIESRHRQESGFSCKGPGQNVAVTRAHPDSQGRRRRPERGARGGQVFYNSEYGELSEPSEEDHCSPSARVTFFTDNSY |
| **Type 5 RTKs (fibroblast growth factor receptor) family** | |
| FGFR1: fibroblast growth factor receptor 1 | DLSMPLDQYSPSFPDTRSSTCSSGEDSVFSHEPLPEEPCLPRHPAQLANGGLKRR |
| FGFR2: fibroblast growth factor receptor 2 | DLSQPLEQYSPSYPDTRSSCSSGDDSVFSPDPMPYEPCLPQYPHINGSVKT |
| FGFR3: fibroblast growth factor receptor 3 | DLSAPFEQYSPGGQDTPSSSSSGDDSVFAHDLLPPAPPSSGGSRT |
| FGFR4: fibroblast growth factor receptor 4 | DLRLTFGPYSPSGGDASSTCSSSDSVFSHDPLPLGSSSFPFGSGVQT |
| **Type 6 RTKs (PTK7/CCK4) family** | |
| CCK4: protein tyrosine kinase 7 | DKSP |
| **Type 7 RTKs (neutrophin receptor/Trk) family** | |
| trkA: neurotrophic receptor tyrosine kinase 1 | QALAQAPPVYLDVLG |
| trkB: neurotrophic receptor tyrosine kinase 2 | QNLAKASPVYLDILG |
| trkC: neurotrophic receptor tyrosine kinase 3 | Nil C-terminal to kinase domain |
| **Type 8 RTKs (ROR) family** | |
| ROR1: receptor tyrosine kinase like orphan receptor 1 | RSWEGLSSHTSSTTPSGGNATTQTTSLSASPVSNLSNPRYPNYMFPSQGITPQGQIAGFIGPPIPQNQRFIPINGYPIPPGYAAFPAAHYQPTGPPRVIQHCPPPKSRSPSSASGSTSTGHVTSLPSSGSNQEANIPLLPHMSIPNHPGGMGITVFGNKSQKPYKIDSKQASLLGDANIHGHTESMISAEL |
| ROR2: receptor tyrosine kinase like orphan receptor 2 | RAWGNLSNYNSSAQTSGASNTTQTSSLSTSPVSNVSNARYVGPKQKAPPFPQPQFIPMKGQIRPMVPPPQLYVPVNGYQPVPAYGAYLPNFYPVQIPMQMAPQQVPPQMVPKPSSHHSGSGSTSTGYVTTAPSNTSMADRAALLSEGADDTQNAPEDGAQSTVQEAEEEEEGSVPETELLGDCDTLQVDEAQVQLEA |
| **Type 9 RTKs (MuSK) family** | |
| MuSK: muscle-associated receptor tyrosine kinase | ERMCERAEGTVSV |
| **Type 10 RTKs (hepatocyte growth factor receptor) family** | |
| MET: MET proto-oncogene RTK | GEHYVHVNATYVNVKCVAPYPSLLSSEDNADDEVDTRPASFWETS |
| Ron: macrophage stimulating 1 receptor | SALLGDHYVQLPATYMNLGPSTSHEMNVRPEQPQFSPMPGNVRRPRPLSEPPRPT |
| **Type 11 RTKs (TAM) family** | |
| Axl: AXL RTK | KALPPAQEPDEILYVNMDEGGGYPEPPGAAGGADPPTQPDPKDSCSCLTAAEVHPAGRYVLCPSTTPSPAQPADRGSPAAPGQEDGA |
| Tyro3: TYRO3 RTK | GQLSVLSASQDPLYINIERAEEPTAGGSLELPGRDQPYSGAGDGSGMGAVGGTPSDCRYILTPGGLAEQPGQAEHQPESPLNETQRLLLLQQGLLPHSSC |
| Mer: MER proto-oncogene, tyrosine kinase | ESLPDVRNQADVIYVNTQLLESSEGLAQGSTLAPLDLNIDPDSIIASCTPRAAISVVTAEVHDSKPHEGRYILNGGSEEWEDLTSAPSAAVTAEKNSVLPGERLVRNGVSWSHSSMLPLGSSLPDELLFADDSSEGSEVLM |
| **Type 12 RTKs (TIE angiopoietin receptor) family** | |
| TIE1: tyrosine kinase with Ig-like and EGF-like domains | NMSLFENFTYAGIDATAEEA |
| TIE2: TEK RTK | EERKTYVNTTLYEKFTYAGIDCSAEEAA |
| **Type 13 RTKs (ephrin receptor) family** | |
| EphA1: EPH receptor A1 | SAM C-terminal to kinase domain, then ends |
| EphA2: EPH receptor A2 | SAM C-terminal to kinase domain, then VNTVGIPI |
| EphA3: EPH receptor A3 | SAM C-terminal to kinase domain, then SKNGPVPV |
| EphA4: EPH receptor A4 | SAM C-terminal to kinase domain, then MQQMHGRMVPV |
| EphA5: EPH receptor A5 | SAM C-terminal to kinase domain, then LVNGMVPL |
| EphA6: EPH receptor A6 | SAM C-terminal to kinase domain, then MMHIQEKGFHV |
| EphA7: EPH receptor A7 | SAM C-terminal to kinase domain, then MLHLHGTGIQV |
| EphA8: EPH receptor A8 | SAM C-terminal to kinase domain, then LTSTQG |
| EphA10: EPH receptor A10 | SAM C-terminal to kinase domain, then VLQ |
| EphB1: EPH receptor B1 | SAM C-terminal to kinase domain, then ISQSP |
| EphB2: EPH receptor B2 | SAM C-terminal to kinase domain, then MNQIQSVEGQPLARRPRATGRTKRCQPRDVTKKTCNSNDGKKKGMGKKKTDPGRGREIQGIFFKEDSHKESNDCSCGG |
| EphB3: EPH receptor B3 | SAM C-terminal to kinase domain, then MNQTLPVQV |
| EphB4: EPH receptor B4 | SAM C-terminal to kinase domain, then AKPGTPGGTGGPAPQY |
| EphB6: EPH receptor B6 | SAM C-terminal to kinase domain, then LRQQGSVE |
| **Type 14 RTKs (RET) family** | |
| Ret: ret proto-oncogene | DLAASTPSDSLIYDDGLSEEETPLVDCNNAPLPRALPSTWIENKLYGMSDPNWPGESPVPLTRADGTNTGFPRYPNDSVYANWMLSPSAAKLMDTFDS |
| **Type 15 RTKs (RYK) family** | |
| RYK (receptor-like tyrosine kinase) | GAYV |
| Type 16 RTKs (DDR, collagen receptor) family | |
| DDR1: discoidin domain RTK 1 | AEDALNTV |
| DDR2: discoidin domain RTK 2 | LQQGDE |
| **Type 17 RTKs (ROS) family** | |
| ROS: c-ros oncogene 1 RTK | LNSIYKSRDEANNSGVINESFEGEDGDVICLNSDDIMPVALMETKNREGLNYMVLATECGQGEEKSEGPLGSQESESCGLRKEEKEPHADKDFCQEKQVAYCPSGKPEGLNYACLTHSGYGDGSD |
| **Type 18 RTKs (LMR) family** | |
| Lmr1: apoptosis-associated tyrosine kinase | Proline rich region at 709-829 |
| Lmr2: lemur tyrosine kinase 2 | Proline rich region at 1451-1454 |
| Lmr3: lemur tyrosine kinase 3 | Proline rich region at 416-1457 |
| **Type 19 RTKs (leucocyte tyrosine kinase) family** | |
| LTK: leucocyte RTK | LNSLLPMELGPTPEEEGTSGLGNRSLECLRPPQPQELSPEKLKSWGGSPLGPWLSSGLKPLKSRGLQPQNLWNPTYRS |
| ALK: ALK RTK | INTALPIEYGPLVEEEEKVPVRPKDPEGVPPLLVSQQAKREEERSPAAPPPLPTTSSGKAAKKPTAAEISVRVPRGPAVEGGHVNMAFSQSNPPSELHKVHGSRNKPTSLWNPTYGSWFTEKPTKKNNPIAKKEPHDRGNLGLEGSCTVPPNVATGRLPGASLLLEPSSLTANMKEVPLFRLRHFPCGNVNYGYQQQGLPLEAATAPGAGHYEDTILKSKNSMNQPGP |
| **Type 20 RTKs (STYK1) family** | |
| STYK1: serine/threonine/tyrosine kinase 1 | KTADDEAVLQVPELVVPELYAAVAGIRVESLFYNYSML |
