## Supplementary figures and images for "Affinity purification mass spectrometry characterization of the interactome of receptor tyrosine kinase proline rich motifs in cancer"

### Supplementary Figures

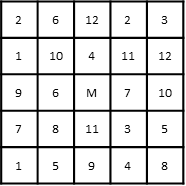

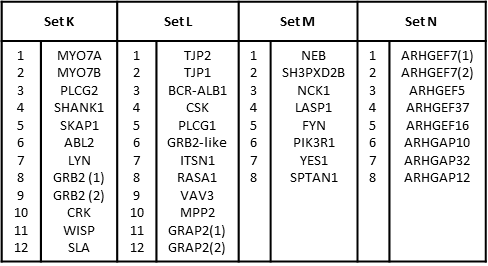

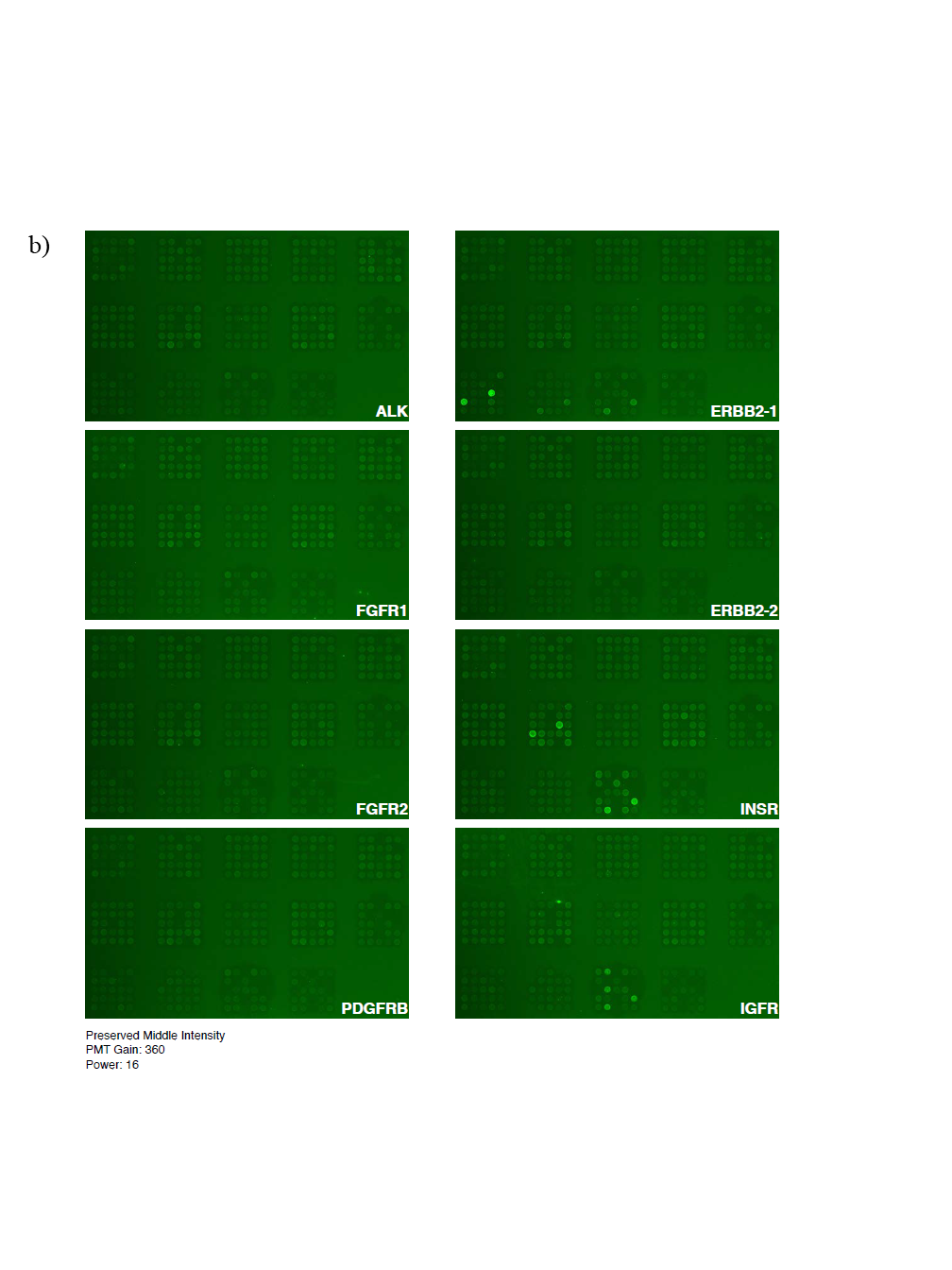

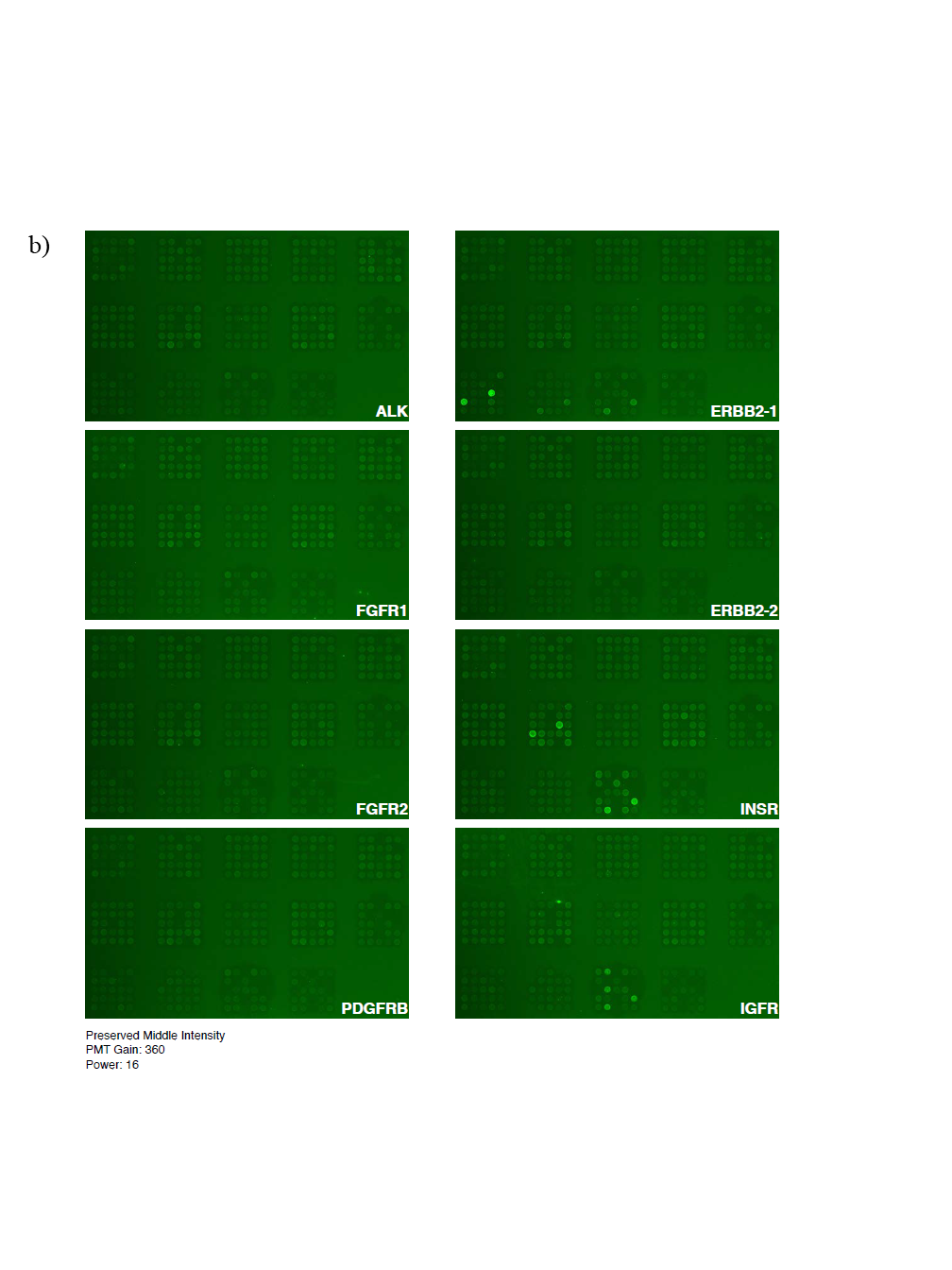

**Supplementary Figure 1**

Set K

Set L

Set M

Set N

**Supplementary Figure 2**

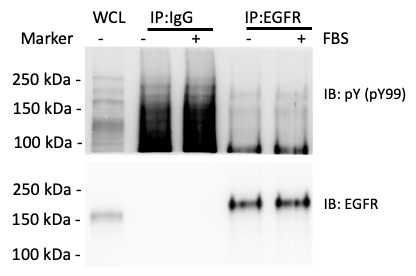

**Supplementary Figure 3**

a)

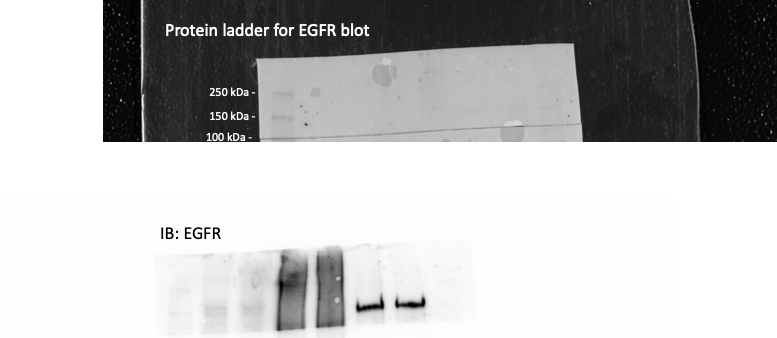

b)

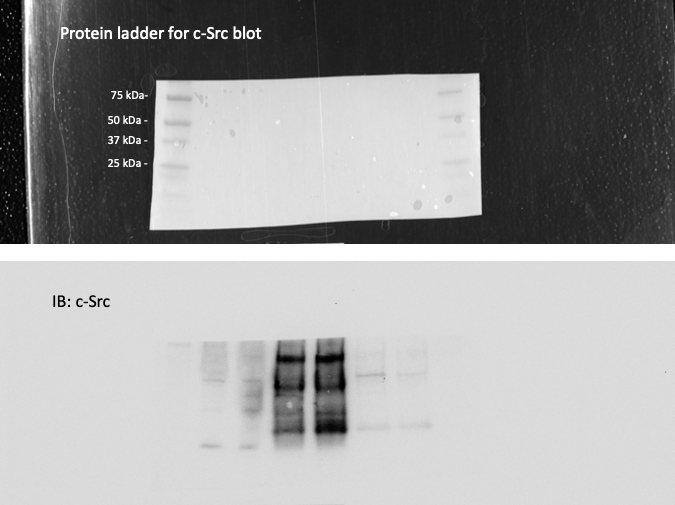

c)

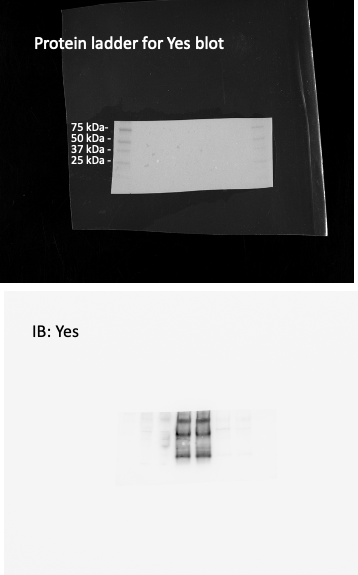

d)
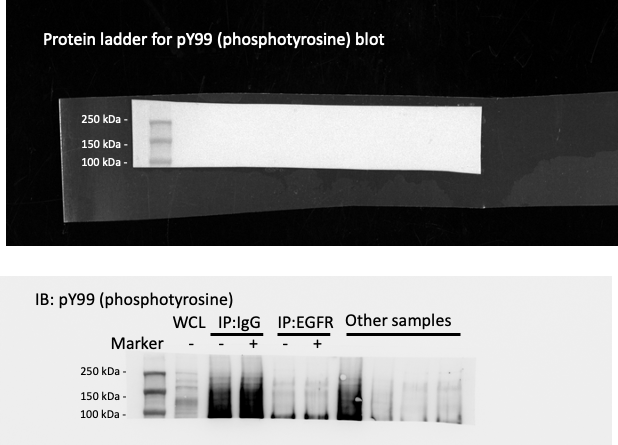
