## Supplementary Material Legends for "Affinity purification mass spectrometry characterization of the interactome of receptor tyrosine kinase proline rich motifs in cancer"

**SUPPLEMENTARY FIGURE LEGENDS**

**Supplementary Figure 1: Control peptide microarray.** A peptide microarray demonstrating interactions between the c-terminal tail of IGF1R, which does not incorporate a proline-rich motif, and glutathione S-transferase (GST) fusion SH3-domain containing proteins spotted on to a positional grid. The correlation between each protein and its grid position is shown alongside. Bound peptides are those with two fluorescent spots present. The experimental peptide microarray is shown in **Fig. 4a.**

**Supplementary Figure 2: Tyrosine phosphorylation on EGFR is not upregulated upon FBS exposure in 293T cells.** EGFR was immunoprecipitated in 293T cells that were either starved or stimulated with FBS. It was then assessed for phosphorylation level in its tyrosine residues using anti-phosphotyrosine antibody (pY99). No evidence for additional phosphorylation is observed in the presence of FBS.

**Supplementary Figure 3: Uncropped Western blot images for (a) EGFR, (b) SRC, (c) YES and (d) phosphotyrosine residues.**

**SUPPLEMENTARY TABLE LEGENDS**

**Supplementary Table 1: Proline-rich motifs and other, less common, Src homology 3 (SH3) domain-binding sequences within the cytoplasmic tails of all 58 receptor tyrosine kinases (RTKs) encoded by the human genome.** Common SH3 domain-binding RTK tail sequences (PxxP) are highlighted in yellow. Cannonical PxxPxR, RxPxxP and RxxPxxP motifs are underlined. Less common SH3 domain-binding RTK tail sequences are shown in purple (PxxDY), blue (PxPxxR) and green (R/KxLP).

**Supplementary Table 2: Interactors, listed by receptor tyrosine kinase and cell line.** Interacting proteins shown were identified using the Significance Analysis of INTeractome (SAINT) scoring algorithm.

**Supplementary table 3: Significance Analysis of INTeractome (SAINT) analysis of epidermal growth factor receptor (EGFR) recovered by affinity purification mass spectrometry in lung squamous cell carcinoma (H520) cells.**

**Supplementary table 4: Significance Analysis of INTeractome (SAINT) analysis of fibroblast growth factor receptor 2 (FGFR2) recovered by affinity purification mass spectrometry in lung squamous cell carcinoma (H520) cells.**

**Supplementary table 5: Significance Analysis of INTeractome (SAINT) analysis of erb-B2 receptor tyrosine kinase (ERBB2/HER2) recovered by affinity purification mass spectrometry in lung squamous cell carcinoma (H520) cells.**

**Supplementary table 6: Significance Analysis of INTeractome (SAINT) analysis of epidermal growth factor receptor (EGFR) recovered by affinity purification mass spectrometry in human embryonic kidney (HEK)-293 cells.**

**Supplementary table 7: Significance Analysis of INTeractome (SAINT) analysis of fibroblast growth factor receptor 2 (FGFR2) recovered by affinity purification mass spectrometry in human embryonic kidney (HEK)-293 cells.**

**Supplementary table 8: Significance Analysis of INTeractome (SAINT) analysis of fibroblast growth factor receptor 2 (FGFR2) recovered by affinity purification mass spectrometry in human embryonic kidney (HEK)-293 cells.**

**Supplementary table 9: Significance Analysis of INTeractome (SAINT) analysis of epidermal growth factor receptor (EGFR) recovered by affinity purification mass spectrometry in oesophageal adenocarcinoma (OE19) cells.**

**Supplementary table 10: Significance Analysis of INTeractome (SAINT) analysis of fibroblast growth factor receptor 2 (FGFR2) recovered by affinity purification mass spectrometry in oesophageal adenocarcinoma (OE19) cells.**

**Supplementary table 11: Significance Analysis of INTeractome (SAINT) analysis of erb-B2 (ERBB2/HER2) recovered by affinity purification mass spectrometry in oesophageal adenocarcinoma (OE19) cells.**

**Supplementary table 12: Significance Analysis of INTeractome (SAINT) analysis of epidermal growth factor receptor (EGFR) recovered by affinity purification mass spectrometry in breast adenocarcinoma (SK-BR-3) cells.**

**Supplementary table 13: Significance Analysis of INTeractome (SAINT) analysis of fibroblast growth factor receptor 2 (FGFR2) recovered by affinity purification mass spectrometry in breast adenocarcinoma (SK-BR-3) cells.**

**Supplementary table 14: Significance Analysis of INTeractome (SAINT) analysis of erb-B2 (ERBB2/HER2) recovered by affinity purification mass spectrometry in breast adenocarcinoma (SK-BR-3) cells.**

**Supplementary table 15: Significance Analysis of INTeractome (SAINT) analysis of epidermal growth factor receptor (EGFR) recovered by affinity purification mass spectrometry in glioblastoma (U251) cells.**

**Supplementary table 16: Significance Analysis of INTeractome (SAINT) analysis of fibroblast growth factor receptor 2 (FGFR2) recovered by affinity purification mass spectrometry in glioblastoma (U251) cells.**

**Supplementary table 17: Significance Analysis of INTeractome (SAINT) analysis of erb-B2 (ERBB2/HER2) recovered by affinity purification mass spectrometry in glioblastoma (U251) cells.**

**Supplementary table 18: pfam domains and their respective clans for each protein identified by affinity purification mass spectrometry.**
